# Detailed temporal dissection of an enhancer cluster reveals two distinct roles for individual elements

**DOI:** 10.1101/2020.05.06.080564

**Authors:** Henry Thomas, Elena Kotova, Axel Pilz, Merrit Romeike, Andreas Lackner, Christoph Bock, Martin Leeb, Joanna Wysocka, Christa Buecker

## Abstract

Many genes are regulated by multiple enhancers that often simultaneously activate their target gene. Yet, how individual enhancers collaborate to activate transcription is not well understood. Here, we dissect the functions and interdependencies of five enhancer elements that form a previously identified enhancer cluster and activate the *Fgf5* locus during exit from naïve murine pluripotency. Four elements are located downstream of the *Fgf5* gene and form a super-enhancer. Each of these elements contributes to *Fgf5* induction at a distinct time point of differentiation. The fifth element is located in the first intron of the *Fgf5* gene and contributes to *Fgf5* expression at every time point by amplifying overall *Fgf5* expression levels. This amplifier element strongly accumulates paused RNA Polymerase II but does not give rise to a mature *Fgf5* mRNA. By transplanting the amplifier to a different genomic position, we demonstrate that it enriches for high levels of paused RNA Polymerase II autonomously. Based on our data, we propose a model for a mechanism by which RNA Polymerase II accumulation at a novel type of enhancer element, the amplifier, contributes to enhancer collaboration.

## Introduction

During development, changes in gene expression are tightly controlled, to allow for the embryo to undergo numerous cell fate transitions. Cis-regulatory elements such as enhancers determine when and how genes are activated. Enhancers are short stretches of DNA consisting of multiple transcription factor binding sites that are located within the non-coding part of the genome and activate transcription of their target gene from a distance (Catarino & Stark, 2018; Long *et al*., 2016). Upon activation of enhancers, transcription factors bind, facilitate removal of nucleosomes and recruit co-activators such as p300. This leads to specific histone modifications on the surrounding nucleosomes such as H3K27ac and H3K4me1 (Catarino & Stark, 2018; Long *et al*., 2016; Visel *et al*., 2009). In addition, RNA Polymerase II (Pol II) itself is also recruited to enhancers, which results in transcription of short-lived RNAs referred to as enhancer RNAs (eRNAs) (Kim *et al*., 2010; Schwalb *et al*., 2016).

Active enhancers in a cell type of interest can be identified based on enhancer-specific chromatin features such as accessible chromatin, p300 binding and accumulation of H3K27ac (Long *et al*., 2016). Such studies have been carried out in numerous cell lines and tissues, to map the regulatory landscape during development and in cancer (Long *et al*., 2016). They also identified so-called stretch- or super-enhancers (SEs) (Parker *et al*., 2013; Whyte *et al*., 2013): clusters of enhancers spanning multiple kilobases (kb) of genomic DNA that are active in the same cell type and collaborate to regulate their target gene (Hnisz *et al*., 2013). SEs are characterized by particularly strong accumulation of the mediator complex, Pol II, p300 and histone modifications such as H3K27ac (Hnisz *et al*., 2013; Whyte *et al*., 2013).

Often, target genes of SEs are highly expressed and of particular importance for the cell type of interest. However, previous studies have provided conflicting results on whether SEs are indeed different from regular enhancers (Moorthy *et al*., 2017), and on the importance of individual elements within these enhancer clusters (Hay *et al*., 2016; Shin *et al*., 2016). At some loci, each element contributes additively and independently to the overall output from the promoter without obvious higher-order effects (Hay *et al*., 2016). At other loci, some elements were shown to be more important than others, and these elements - referred to in some studies as hub enhancers - might in fact control the activation of other enhancer elements within the same enhancer cluster (Hnisz *et al*., 2015, Huang *et al*., 2018; Shin *et al*., 2016; Xu *et al*., 2012). Finally, for the *Fgf8* locus, both scenarios have been observed recently, with *Fgf8* expression being regulated either by a set of redundant enhancers or by a combination of one dominant enhancer and two enhancers with only minor impact, depending on the analyzed cell type (Hörnblad *et al*., 2020).

Since most target genes of SEs are vital for their specific cell state (Whyte *et al*., 2013), any perturbation leading to lower expression of the target gene could in turn affect this particular cell state. Therefore, conclusions about the detailed contributions of individual elements to transcription of their target genes must be very carefully disentangled from changes in cell state that might in turn feedback on target gene expression. Furthermore, how an enhancer cluster is activated during transition from one cell state to a closely related one remains unclear, since enhancer clusters have mostly been studied at a defined stage of development. Are all enhancer elements activated at the same time and contribute to expression at all time points of a cell fate transition, or do different enhancer elements affect distinct time points?

In this study, we dissected the contributions of individual enhancer elements constituting an enhancer cluster to the activation of their target gene during the transition from one cell state to a closely related one. We took advantage of the well-characterized changes within the enhancer landscape during the exit from naïve pluripotency. We have previously identified the *Fgf5* enhancer cluster that is activated during the exit from naïve pluripotency (Buecker *et al*., 2014). As *Fgf5* is dispensable for early embryonic development, this enhancer cluster provides a good model for studying enhancer collaboration. Through careful temporal dissection, we show here that the enhancer elements at the *Fgf5* locus fall into two classes of regulatory elements: While the intergenic enhancers E1-E4 contribute to induction of *Fgf5* expression at specific time points of exit from naïve pluripotency, the intronic PE enhancer amplifies expression levels at all time points. All five elements are required to achieve full expression of the target gene, and PE collaborates with the enhancers E1-E4 in a super-additive fashion. Finally, we observed high levels of Pol II at PE, and we suggest that PE works as an amplifier element by increasing the local concentration of Pol II, thus boosting overall expression levels at the *Fgf5* locus.

## Results

### Identification of the *Fgf5* enhancer cluster as a model locus for collaborative enhancer action

Dissecting the temporal contributions of individual enhancer elements within an SE can be hampered by the importance of the target gene for correctly establishing the cell state of interest: if deletion of individual enhancers lowers transcription of the target gene, this decrease in target gene expression might in turn change the overall cell state. Direct consequences of enhancer deletions on target gene expression are therefore difficult to disentangle from indirect ones arising from a change in cell state, such as different expression levels of transcription factors and co-activators. We have previously characterized the changes in the enhancer landscape during the transition from naïve pluripotent mouse embryonic stem cells (ESCs) into the closely related cell state, epiblast like cells (EpiLCs). The transition is often also referred to as the exit from naïve pluripotency. We have identified the *Fgf5* enhancer cluster as a model system to study the interaction among individual enhancer elements during cell fate transition in detail (Buecker *et al*., 2014). The *Fgf5* enhancer cluster consists of five individual elements: E1 through E4 are located between 29 and 58 kb downstream of the transcription start site (TSS) within the non-coding part of the genome (Fig 1A). These four elements together form a SE as defined by the ROSE-algorithm (Whyte *et al*., 2013), based on H3K27ac deposition and p300 accumulation at neighboring elements with a maximum distance of 12.5 kb (Fig S1A). These putative enhancer elements are in an off-state in ESCs, with no detectable enhancer marks and closed chromatin. During differentiation into EpiLCs, all sites gain H3K27ac, H3K4me1, p300, and OCT4 (Fig 1A, and data not shown). In addition, a fifth putative enhancer element is located within the first intron of *Fgf5* less than two kb from the TSS. This element is already accessible in the ESC state and pre-bound by low levels of p300 and OCT4 (Fig 1A and data not shown), however, H3K27ac is deposited only during differentiation. Instead, the promoter and the enhancer are marked by low levels of the repressive H3K27me3 mark in the ESC state that are removed upon differentiation (Fig 1A). We therefore refer to this element as poised enhancer (PE) (Rada-Iglesias *et al*., 2011).

**Figure 1:**
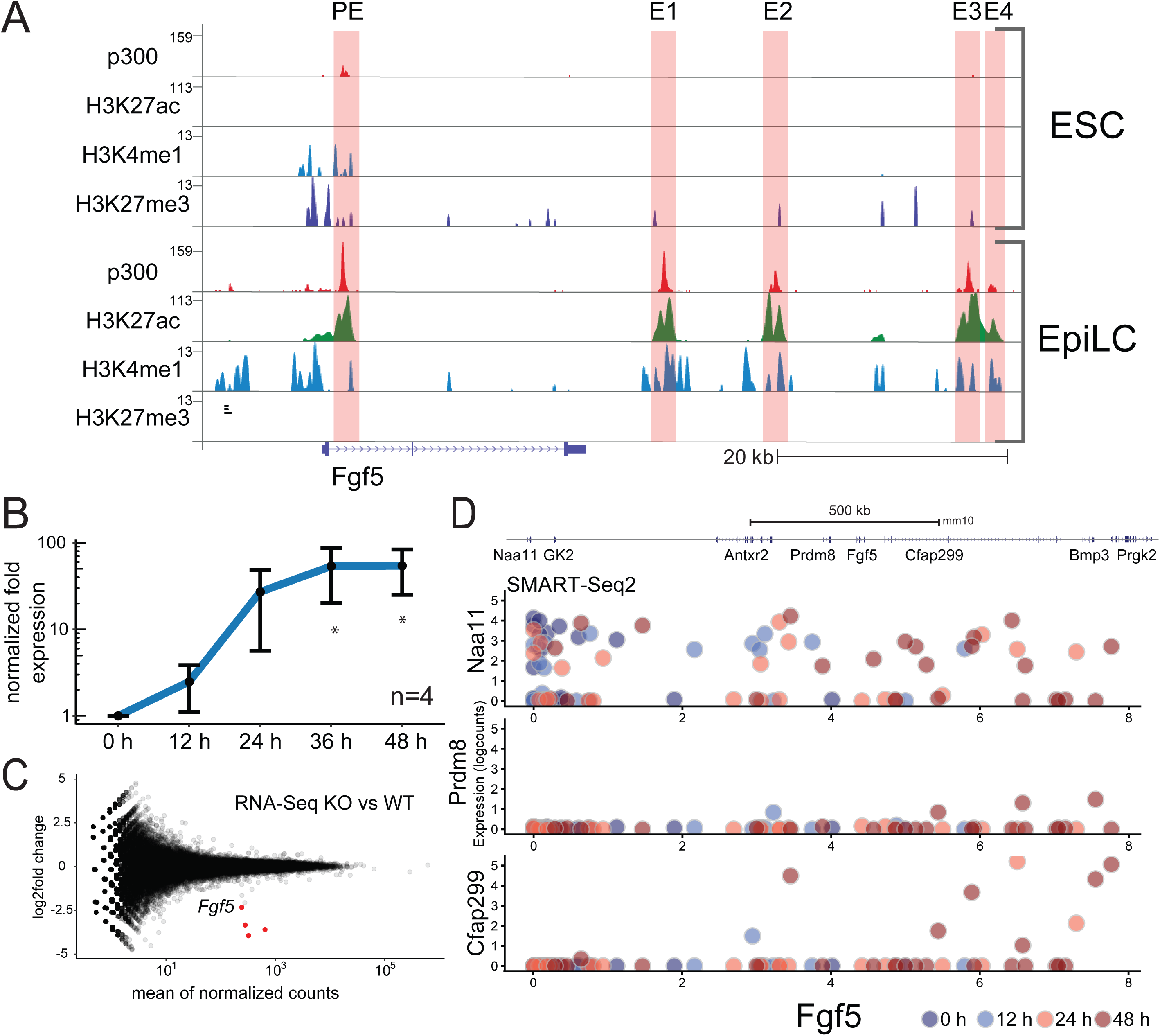
The *Fgf5* locus as a model to study collaborative enhancer action. (A) ChIP-Seq signal for p300, H3K27ac, H3K4me1 and H3K27me3 at the *Fgf5* locus in WT ESCs and EpiLCs from Buecker *et al*., 2014. Putative enhancer elements based on H3K4me1, H3K27ac and p300 ChIP-Seq signal are highlighted with red boxes. (B) *Fgf5* expression in WT cells along an ESC to EpiLC differentiation time course as determined by RT-qPCR with intron-spanning primers. Expression values are normalized to *Rpl13*a and to the 0 h time point within each independent biological replicate. Mean values of n=4 biological replicates are shown. Error bars correspond to one standard deviation in each direction. Time points with statistically significant higher expression (one-sided Welch Two sample t-test) compared to 0 h are marked by stars. (C) Differential expression analysis of WT vs PE KO cell line at 48 h of differentiation. Differential expression analysis on RiboZero RNA-Seq data of two biological replicates each was performed with DESeq2 (Love *et al*., 2014). Differentially expressed genes (log2fold change ≥ 1, adjusted p-value ≤ 0.05) are marked in red. (D) SMART-Seq2 single-cell expression data of genes surrounding *Fgf5* at 0, 12, 24 and 48 h of ESC to EpiLC differentiation. Normalized log counts of the respective gene are plotted against normalized log counts of *Fgf5* in the same cell.

*Fgf5* expression is induced during differentiation in a highly reproducible fashion: the expression within the differentiating population increases steadily to reach a maximum around 36-48 hours (h) after medium exchange (Fig 1B). The expression of *Pou5f1*/*Oct4* does not change during this time frame (data not shown). In contrast, known markers for the EpiLC state such as *Otx2* and *Pou3f1*/*Oct6* are upregulated, whereas naïve pluripotency markers such as *Tbx3* are downregulated (Fig S1B and data not shown).

Importantly, while *Fgf5* negatively controls hair growth later in development, it is dispensable for early embryonic development (Hébert *et al*., 1994). This makes it an excellent model locus for genetic enhancer studies, as perturbing *Fgf5* expression levels does not affect differentiation *per se*. To confirm that *Fgf5* is indeed dispensable for the exit from naïve pluripotency, we performed RiboZero RNA-Seq in wild type (WT) cells at 48 h of differentiation and compared the results to an enhancer knock-out (KO) cell line that shows a 10-fold decrease in *Fgf5* expression levels. Despite drastically reduced *Fgf5* levels, we did not observe major changes in overall gene expression, as only three genes (*Egr1*, *Eif2s3y*, *Uty*) besides *Fgf5* showed statistically significant changes (Fig 1C).

*Fgf5* is located in a small topologically associated domain (TAD) on chromosome five along with either *Prdm8* (Hi-C data from Rao *et al*., 2014) or together with *Prdm8*, *Cfap299* and *Bmp3* (Hi-C data from Dixon *et al*., 2012). We tested whether any of the surrounding genes might be regulated by the enhancers at the *Fgf5* locus. As these enhancers are only activated upon exit from pluripotency (Fig 1A), we expect such genes to be upregulated during differentiation. We performed SMART-Seq2 single cell RNA-Seq along a time course with high temporal resolution to account for the intrinsic heterogeneity of the differentiation process (Chaigne *et al*., 2019). *Fgf5* was upregulated in the majority of cells during differentiation (Fig 1D), and can thus serve as a marker for progression of differentiation. We compared the expression of *Fgf5* against the expression of each of the surrounding genes within two megabases (MB) in the exact same cell throughout differentiation, as expression of genes upregulated during differentiation should correlate with *Fgf5* expression. *Prdm8* was slightly upregulated in very few cells, whereas *Cfap299* expression was strongly upregulated, but only in few cells (Fig 1D). The only other expressed gene within one MB of the *Fgf5* TSS was *Naa11* (Fig 1D and S1C), however, expression of *Naa11* did not change during differentiation and was not correlated with *Fgf5* expression. In addition, none of the surrounding genes were differentially expressed in the RNA-Seq comparison between WT and KO cell line (data not shown). This indicates that the enhancer elements at the locus indeed regulate *Fgf5*, rather than the surrounding genes.

Taken together, *Fgf5* is strongly induced during the ESC to EpiLC transition, but reduced *Fgf5* levels do not perturb the differentiation process. Due to this absence of potential indirect effects and its genomic location with few surrounding genes being expressed, we conclude that the *Fgf5* enhancer cluster is a suitable model locus to dissect the contributions of individual enhancer elements to target gene expression with high temporal resolution along the transition from one cell type to a closely related one.

### Individual SE elements contribute to *Fgf5* induction at distinct time points

To study the effect of putative enhancer elements on *Fgf5* expression, we deleted individual enhancers using CRISPR/Cas9. Therefore, we designed single guide RNAs flanking the p300 peak and isolated clones carrying homozygous deletions of the targeted enhancer element. For each enhancer KO, we tested several independent clones with similar results. We also confirmed that the ESC to EpiLC differentiation is not affected due to clonal effects by testing the expression changes of known ESC and EpiLC markers (*Tbx3*, *Rex1* and *Pou3f1/Oct6*, *Otx2,* respectively; data not shown). We differentiated KO ESC lines to EpiLCs and quantified *Fgf5* expression levels by RT-qPCR at different time points. While we did observe consistent trends for the different KO cell lines compared to WT, overall expression levels varied between biological replicates due to the variability associated with the differentiation process (as can be seen in Fig 1B for WT). Therefore, to assess the significance of our observations, we decided for the following strategy to present and normalize our data. Average expression values were calculated based on several biological replicates for each cell line, and are shown as line graphs along the ESC to EpiLC differentiation. These line graphs give an overview of how the different cell lines behave compared to WT and are shown without error bars (e. g. Fig 2A). For quantitative comparisons, we normalized the expression value of each cell line and time point to the expression value of a WT cell line that has been differentiated in parallel. These WT-normalized values are depicted in bar graphs and are used to determine significantly different expression values as compared to WT at individual time points (e. g. Fig 2B).

**Figure 2:**
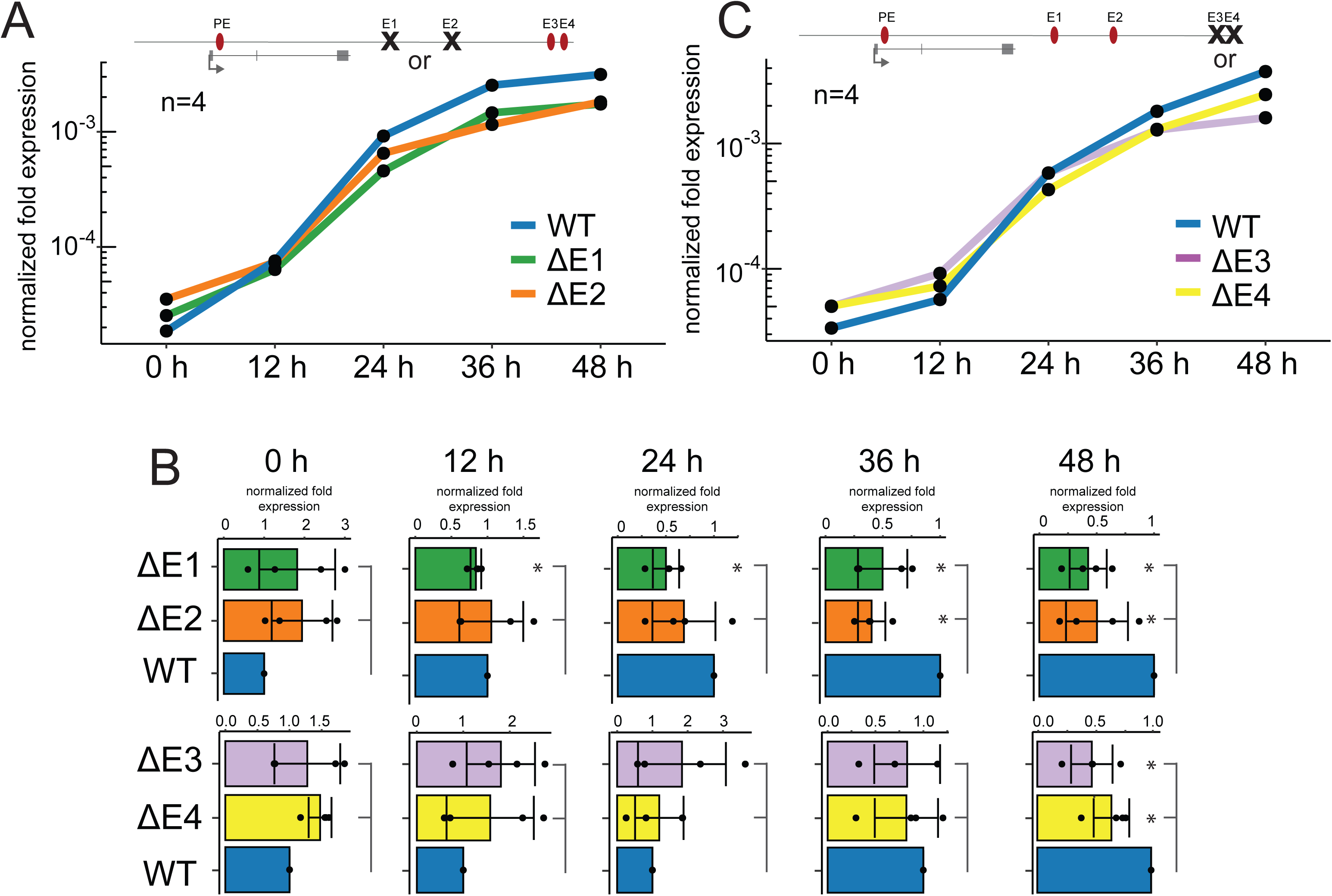
The intergenic enhancers E1-E4 mediate induction of *Fgf5* upon differentiation. (A) *Fgf5* expression in WT, ΔE1 and ΔE2 cells along an ESC to EpiLC differentiation time course as determined by RT-qPCR with intron-spanning primers. Expression values are normalized to *Rpl13a*. Mean values of n=4 biological replicates are shown. (B) *Fgf5* expression in WT, ΔE1, ΔE2, ΔE3 and ΔE4 cells at each time point of ESC to EpiLC differentiation as determined by RT-qPCR with intron-spanning primers. Expression values are normalized to *Rpl13a* and to the WT cell line at the same time point within each biological replicate. Mean values of n=4 biological replicates are shown. Normalized values for each replicate are shown as dots. Error bars correspond to one standard deviation in each direction. Cell lines with statistically lower expression (one-sided Welch Two sample t-test) compared to WT at that time point are marked by stars. (C) *Fgf5* expression in WT, ΔE3 and ΔE4 cells along an ESC to EpiLC differentiation time course as determined by RT-qPCR with intron-spanning primers. Expression values are normalized to *Rpl13a*. Mean values of n=4 biological replicates are shown.

The SE of *Fgf5* consists of the four putative enhancer elements E1 through E4. Individual deletion of E1 or E2 did not significantly affect *Fgf5* expression levels in undifferentiated cells (Fig. 2A and 2B), however, upon differentiation, expression of *Fgf5* in these cell lines did not reach WT levels. This reduction of expression levels compared to WT was especially apparent at 36 and 48 h of differentiation, although a significant but very small reduction in the ΔE1 cell line was already observed from 12 h of differentiation forward. Expression of the pluripotency marker *Tbx3* and the differentiation marker *Pou3f1*/*Oct6* were not affected in either cell line (Fig S2A).

Next, we focused on E3 and E4. Similar to E1 and E2, deletion of either element had no significant effect on *Fgf5* expression in undifferentiated ESCs. Upon differentiation, expression levels of *Fgf5* were reduced in the KO cell lines as compared to WT (Fig 2B and 2C). While deletion of E1 and E2 already affected *Fgf5* expression at 36 h of differentiation (or even earlier in the case of E1), E3 and E4 deletion only significantly reduced expression at 48 h, and expression levels in ΔE4 cell lines were slightly higher compared to the other KO cell lines (Fig 2B). Pluripotency and differentiation markers were expressed to similar levels as in WT cells (Fig S2A).

To conclude, the enhancer elements E1-E4 do not contribute to basic levels of *Fgf5* expression in undifferentiated ESCs, but instead mediate the induction of *Fgf5* upon differentiation, with E1 and E2 acting earlier than E3 and E4.

### PE amplifies *Fgf5* expression levels at every time point, yet has little canonical enhancer activity

Deletion of E1 had only minor effects on *Fgf5* expression at 12 h, whereas deletion of E2-E4 did not have any effect (Fig 2B). We therefore asked whether the PE element located within the first intron could be responsible for the early initiation of *Fgf5* expression from 0 h to 12 h. Surprisingly, deletion of this intronic enhancer element reduced *Fgf5* expression at every time point, even in undifferentiated cells (Fig 3A and 3B). In fact, expression levels of *Fgf5* were consistently decreased by roughly 10-fold, leading to a parallel *Fgf5* expression curve that showed the same induction compared to 0 h as in WT cells, but was overall shifted towards lower expression levels. The differentiation process itself was not affected in ΔPE cells (Fig S3A). Therefore, PE seems to “amplify” overall expression levels at the locus at all time points by a factor of 10, whereas E1-E4 specifically induce *Fgf5* expression upon ESC to EpiLC differentiation at distinct time points.

**Figure 3:**
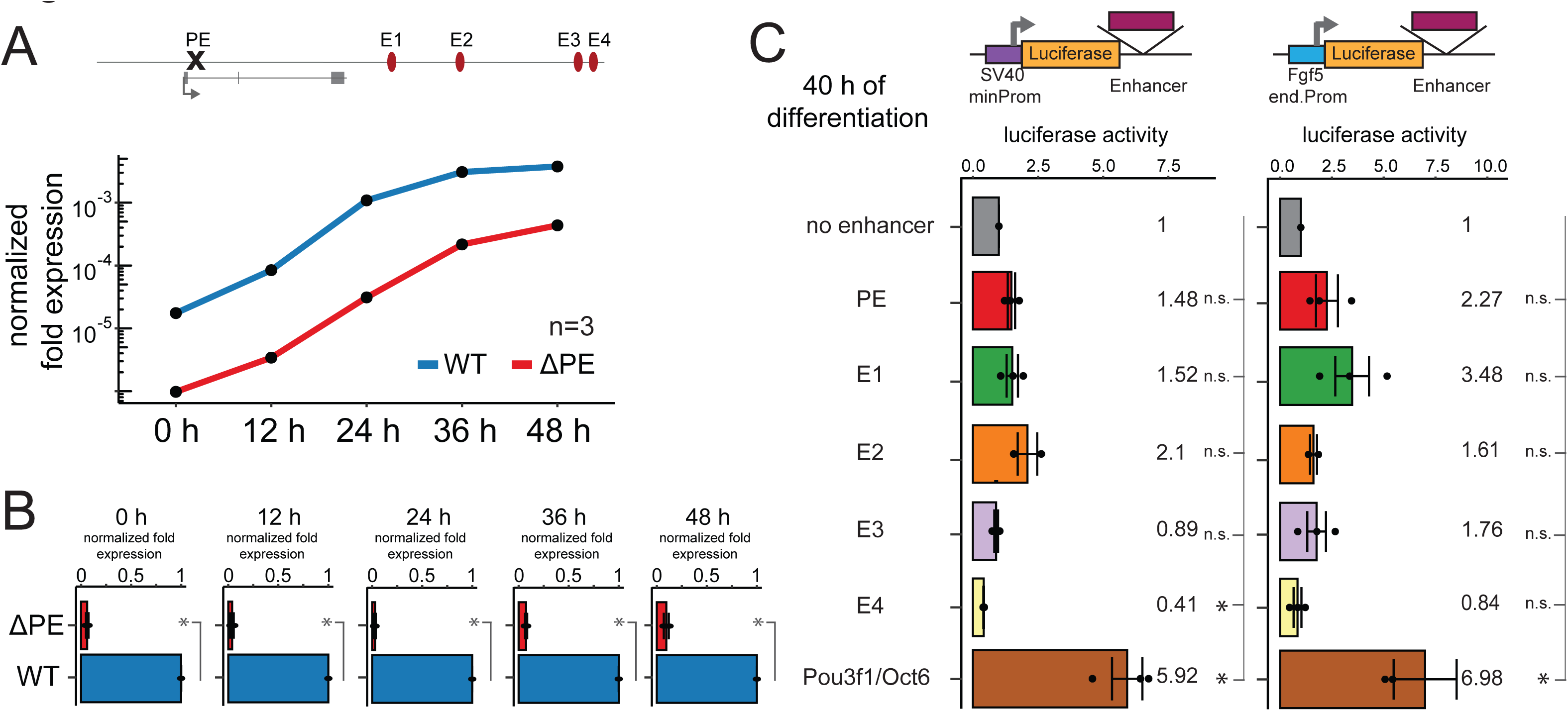
PE amplifies *Fgf5* expression levels at every time point, yet has little canonical enhancer activity in luciferase assays. (A) *Fgf5* expression in WT and ΔPE cells along an ESC to EpiLC differentiation time course as determined by RT-qPCR with intron-spanning primers. Expression values are normalized to *Rpl13a*. Mean values of n=3 biological replicates are shown. (B) *Fgf5* expression in WT and ΔPE cells at each time point of ESC to EpiLC differentiation as determined by RT-qPCR with intron-spanning primers. Expression values are normalized to *Rpl13a* and to the WT cell line at the same time point within each biological replicate. Mean values of n=3 biological replicates are shown. Normalized values for each replicate are shown as dots. Error bars correspond to one standard deviation in each direction. Cell lines with statistically lower expression (one-sided Welch Two sample t-test) compared to WT at that time point are marked by stars. (C) Luciferase assays with the respective enhancer downstream of the luciferase gene under the control of a minimal SV40 promoter (left) or under the control of the endogenous *Fgf5* promoter (right) at 40 h of differentiation. Luciferase activity is normalized first for transfection efficiency as well as plasmid size, and then to the no enhancer control within each independent biological replicate. Mean values of n=3 biological replicates are shown. Normalized values for each replicate are shown as dots. Error bars correspond to one standard deviation in each direction. Enhancers with statistically significant differences (two-sided Welch Two sample t-test) compared to the no enhancer control are marked by stars.

As PE deletion reduced *Fgf5* expression to lower levels than deletion of E1-E4 (Fig 2B and 3B), we tested whether PE also strongly activates transcription in classical assays of enhancer activity. We thus performed luciferase-based enhancer assays. We used two different promoters to ensure enhancer-promoter compatibility, since it has been shown previously that enhancers preferentially activate transcription from certain promoters, while not acting on others (Zabidi *et al*., 2015). We cloned individual enhancers downstream of the luciferase gene under the control of either the SV40 minimal promoter or 495 base pairs (bp) from the endogenous *Fgf5* promoter. We transfected the plasmids into WT ESCs, and started differentiation for 24 or 40 h on the same day. As a positive control, we made use of an enhancer close to the *Pou3f1*/*Oct6* gene that is induced upon differentiation (Fig S2A). This enhancer consistently activated luciferase activity with both promoters at 40 h of differentiation compared to the no enhancer control (Fig 3C). Although deletion of the *Fgf5* enhancers drastically reduced expression at the endogenous locus, none of these enhancers strongly activated luciferase activity at 24 or 40 h of differentiation (Fig 3C and S3B). In fact, none of these constructs showed significantly higher activity than the control plasmid without any enhancers, and E3 and E4 even significantly reduced luciferase activity at some time points (Fig 3C and S3B).

We do note that these luciferase assays were noisy – potentially due to the stress that is put on the cells by starting differentiation a few hours after transfecting the plasmids - and had a limited dynamic range, as even our positive control only induced luciferase activity roughly 6-fold (Fig 3C and S3B). Nonetheless, we were surprised by the low activity of PE in these assays compared to E1-E4 (Fig 3C and S3B), given the much stronger reduction of *Fgf5* expression upon PE deletion (Fig 2B and 3B). In addition, the positive control did activate luciferase activity much more strongly than the *Fgf5* enhancers, demonstrating that despite its limitations the assay is capable of distinguishing stronger from weaker enhancers. Taken together, PE has a strong effect on endogenous expression levels, but only weak canonical enhancer activity in luciferase assays.

### PE collaborates with E1-E4 in a super-additive fashion to regulate transcription of *Fgf5*

Next, we analyzed the expression levels driven by PE in the absence of any additional enhancers at the endogenous *Fgf5* locus. We consecutively deleted all individual elements E1 through E4 and determined the effect on *Fgf5* expression during a differentiation time course. Naïve pluripotency and differentiation markers in this PE only cell line behaved as in WT cells (Fig S4A). While *Fgf5* expression at 0 and 12 h was not significantly affected, expression levels later in differentiation were much reduced compared to WT (Fig 4A and S4B). This confirms that E1-E4 are required for proper induction of *Fgf5* expression during differentiation, while PE acts as an amplifier that determines overall expression levels at the locus. Yet, even in the PE only cell line we detected a small increase in *Fgf5* expression upon differentiation (Fig 4A), therefore we cannot rule out that PE, besides acting as an amplifier, also contributes to induction of *Fgf5* expression.

**Figure 4:**
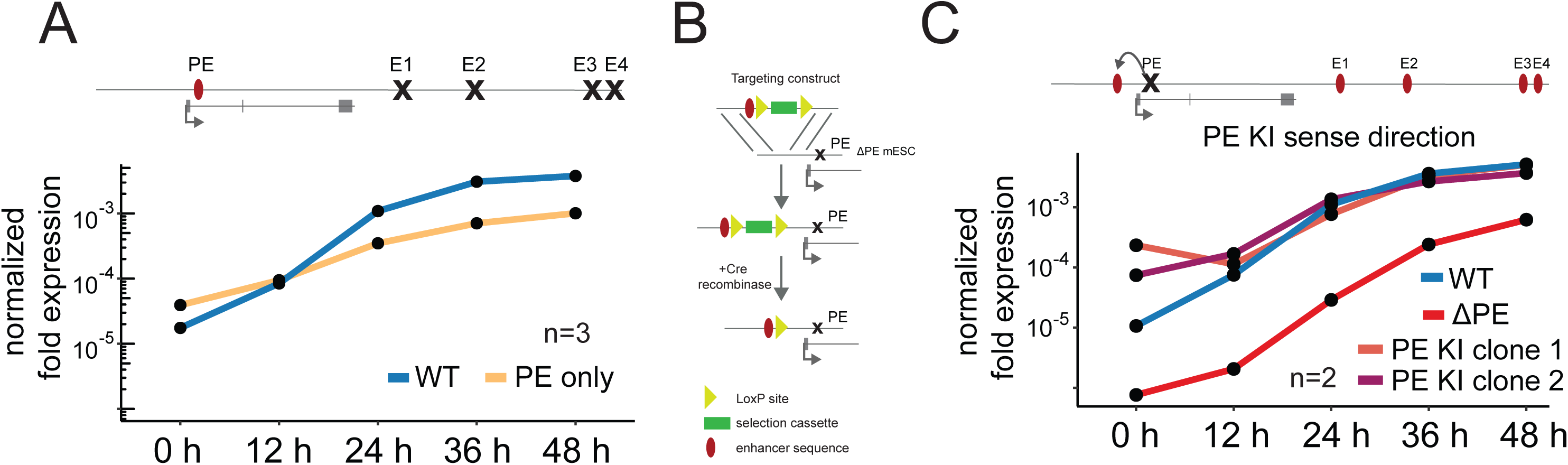
PE and E1-E4 regulate *Fgf5* transcription in super-additive fashion. (A) *Fgf5* expression in WT and PE only (individual deletion of E1 through E4) cells along an ESC to EpiLC differentiation time course as determined by RT-qPCR with intron-spanning primers. Expression values are normalized to *Rpl13a*. Mean values of n=3 biological replicates are shown. (B) Scheme depicting PE KI generation. ΔPE cells were transfected with a linearized targeting construct containing the PE element (red oval) as well as a puro-delta TK selection cassette (green rectangle) flanked by loxP sites (yellow triangles). After integration of this construct upstream of the *Fgf5* promoter, cells were transfected with Cre-recombinase to remove the selection cassette, leaving a single loxP site behind. (C) *Fgf5* expression in WT, ΔPE and PE KI (PE integrated in sense direction) cells along an ESC to EpiLC differentiation time course as determined by RT-qPCR with intron-spanning primers. Expression values are normalized to *Rpl13a*. Mean values of n=2 biological replicates are shown.

Interestingly, deletion of PE reduced expression to around 10% of WT levels, however, in the PE only cell line, *Fgf5* levels amounted to only 25% of WT expression (Fig S4B). This suggests that PE and E1-E4 regulate *Fgf5* expression levels in a super-additive fashion. Under a strictly additive model, one would assume that the expression levels of a PE only cell line – that allows to assess the expression levels driven by PE on its own - and a ΔPE cell line – that allows to assess the expression levels in the absence of PE - added up to 100%. However, this was clearly not the case, as upon differentiation expression levels in ΔPE and PE only cell lines added up to 50% at most (Fig S4B).

Taken together, PE amplifies *Fgf5* expression levels at the endogenous locus at all time points, and collaborates with E1-E4 in a super-additive fashion to achieve WT levels of *Fgf5* expression during differentiation. Yet, despite the greater reduction in expression levels upon deletion at the endogenous locus, canonical enhancer activity of PE in luciferase assays was very low.

We therefore hypothesized that deletion of the intronic sequences in the ΔPE cell line might have disrupted splicing intermediates or RNA modifications that affect RNA production or stability independently of transcriptional regulation (Braunschweig *et al*., 2013; Roundtree *et al*., 2017). To test this, we designed new cell lines in which we re-introduced the PE element into ΔPE cell lines upstream of the *Fgf5* gene (5’ of the TSS), but at a similar distance as in the endogenous location (Fig 4B). We selected multiple clonal cell lines in which PE had been inserted in either sense or antisense direction. After removal of the loxP flanked selection cassette using Cre-recombinase, we measured *Fgf5* expression levels of multiple clones for each orientation during differentiation time courses. In all cases, introduction of the PE element 5’ of the promoter rescued the *Fgf5* expression pattern independently of the direction of the enhancer element, albeit not completely to WT levels (Fig 4C, S4C-F). Interestingly, expression levels at 0 h seemed to be higher in the knock-in (KI) cell lines compared to WT, yet this difference was only significant in one out of four clones and might be caused by higher noise at low expression levels (Fig S4D and S4F). In conclusion, our results suggest that PE regulates transcription rather than splicing as it can exert its function even when not located within the first intron.

### Accumulation of H3K27ac at PE does not occur much earlier compared to E1-E4

PE strongly amplifies *Fgf5* transcription despite low classical enhancer activity in luciferase assays, and affects *Fgf5* expression at every time point, unlike the outside enhancers E1-E4 that are only active later (Fig 2B and 3B). We therefore analyzed the role of PE in activation of *Fgf5* expression in more detail. First, we tested whether earlier activation of PE compared to E1 through E4 could explain the reduced expression levels at very early time points upon PE deletion. We performed ChIP-Seq for H3K27ac along a time course of ESC to EpiLC differentiation. While H3K27ac has been suggested to be dispensable for enhancer function (Bonn *et al*., 2012; Catarino & Stark, 2018; Pengelly *et al*., 2013; Pradeepa *et al*., 2016), deposition of this histone marks strongly correlates with enhancer activity (Bonn *et al*., 2012; Creyghton *et al*., 2010; Heintzman *et al*., 2009; Rada-Iglesias *et al*., 2011; Zentner *et al*., 2011). As expected, accumulation of H3K27ac at the *Pou3f1*/*Oct6* enhancer could be detected as early as 12 h after initiation of differentiation, concomitantly with upregulation of *Pou3f1*/*Oct6* expression, while the *Tbx3* locus lost H3K27ac upon differentiation (Fig S5A). PE did not accumulate H3K27ac immediately after initiation of differentiation, but appreciable amounts of H3K27ac could be detected from 30 h of differentiation on (Fig 5A). At the E3 and E4 enhancers, H3K27ac accumulated at the same time, whereas accumulation at E1 and E2 was only observed at 36 h of differentiation. These findings were corroborated with data from publicly available time courses of ESC differentiation from Yang *et al*., 2019 (Fig S5B), where H3K27ac at PE was detected slightly earlier compared to E1/2 at 24 h, but simultaneously with accumulation at E3 and E4. We conclude that PE, although influencing *Fgf5* expression already in undifferentiated cells, does not accumulate H3K27ac much earlier than E1-E4.

**Figure 5:**
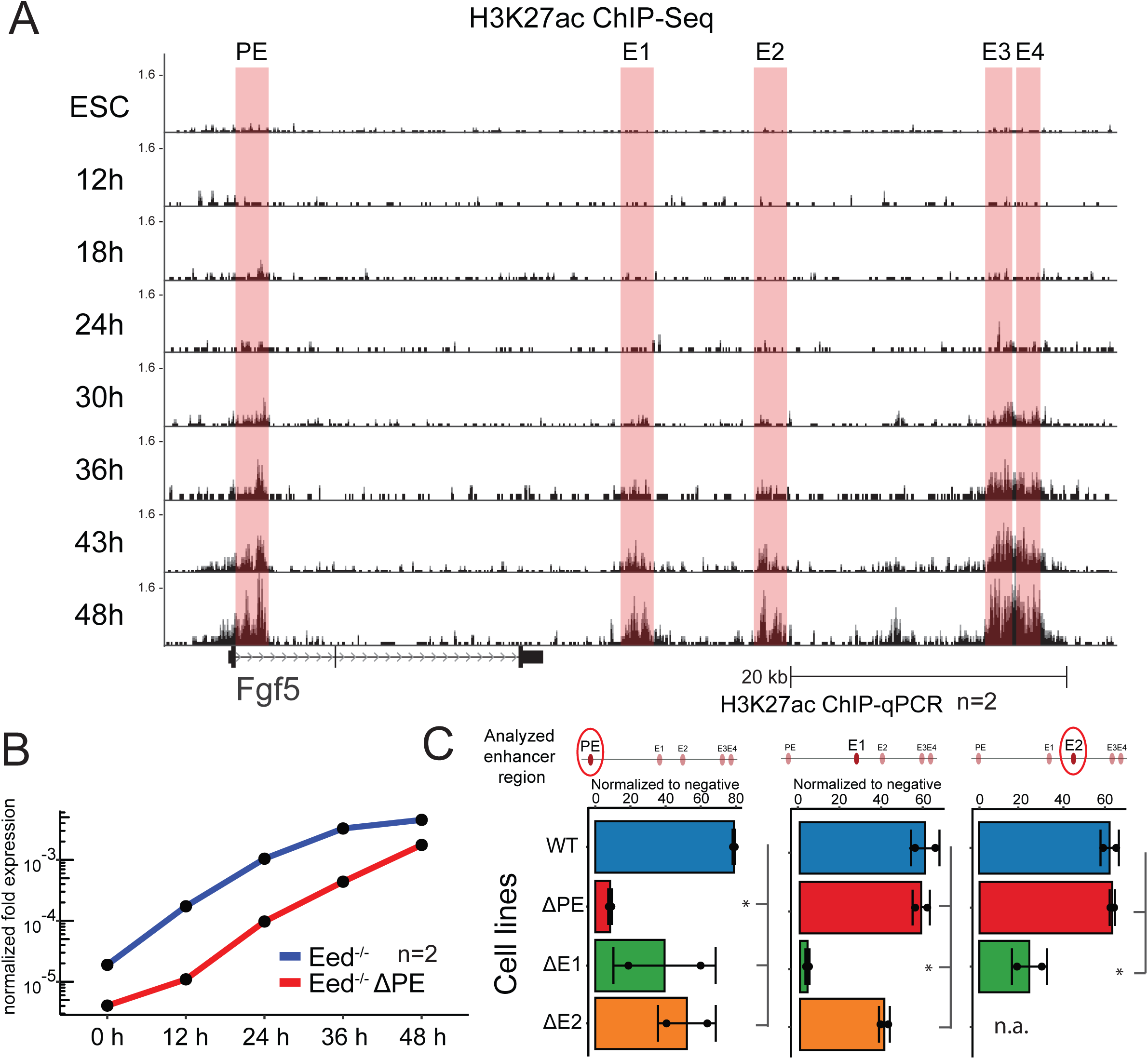
PE is not activated earlier than E1-E4 and does not primarily function by removing H3K27me3 from the *Fgf5* promoter or by facilitating activation of the intergenic enhancers E1-E4. (A) H3K27ac ChIP-Seq signal (normalized for sequencing depth) at the *Fgf5* locus along a differentiation time course with fixed scale bar. (B) *Fgf5* expression in Eed-/- and Eed-/-ΔPE cells along an ESC to EpiLC differentiation time course as determined by RT-qPCR with intron-spanning primers. Expression values are normalized to *Rpl13a*. Mean values of n=2 biological replicates are shown. (C) H3K27ac ChIP-qPCR signal flanking the PE, E1 and E2 enhancers in WT, ΔPE, ΔE1 and ΔE2 cells at 40 h of differentiation. Input enrichment was calculated and then normalized within each individual sample to two genomic regions known not to be marked by H3K27ac by previous ChIP-Seq studies (Buecker *et al*., 2014). Mean values of n=2 biological replicates are shown. Normalized values for each replicate are shown as dots. Error bars correspond to one standard deviation in each direction. Cell lines with statistically lower signal (one-sided Welch Two sample t-test) compared to WT at that time point are marked by stars.

ChIP-Seq and RT-qPCRs are population wide assays that reflect changes across a population of cells, but not within single cells. PE could affect expression in all cells and its deletion could lower *Fgf5* expression across the whole population. Conversely, PE could regulate the probability of *Fgf5* expression, rather than actual expression levels. In this case, *Fgf5* expression would be lost in most cells upon PE deletion, while few single “jackpot” cells would still be able to fully activate *Fgf5* expression. To distinguish between these two scenarios, we performed smRNA-FISH experiments against *Fgf5*, *Otx2* and *Tbx3* using ViewRNA FISH probes (Fig S5C). As expected, Otx2 expression increased across the whole population upon differentiation, and Tbx3 similarly decreased. While neither marker gene was affected by PE deletion, *Fgf5* expression was lower in all ΔPE cells, and we were not able to detect any single cells with high expression of *Fgf5*. We conclude that PE does not regulate probability of *Fgf5* expression, and that it is necessary in all cells to achieve WT expression levels of *Fgf5*.

### PE does not primarily function by counteracting PRC2-mediated H3K27me3 deposition

PE is a poised enhancer, which is marked by both active (p300 and H3K4me1) and repressive (H3K27me3) chromatin marks in undifferentiated cells. During differentiation, the repressive H3K27me3 mark is removed and instead replaced by the active H3K27ac mark (Fig 1A). Upon deletion of PE, *Fgf5* expression is lower and more H3K27me3 can be found surrounding the enhancer, suggesting that the repressive mark is not removed efficiently (data not shown).

We therefore hypothesized that the main function of the PE element could be to counteract H3K27me3 deposition. If that was the case, then global removal of all H3K27me3 should alleviate the need for the PE element. To test this hypothesis, we deleted PE in cells that lack all H3K27me3 due to loss of *Eed.* This gene encodes for a subunit of the PRC2 complex that is responsible for H3K27me3 deposition. *Eed*^-/-^ cells show overall differentiation defects (Lackner *et al*., 2020; Obier *et al*., 2015), however, *Fgf5* expression was strongly upregulated during differentiation (Fig 5B). If the role of PE was only to counteract H3K27me3, then deletion of PE would not affect *Fgf5* expression in cells lacking all K27me3. Yet, we still detected a reduction of *Fgf5* expression upon PE deletion in an *Eed* mutant background at every time point tested (Fig 5B and S5E), whereas pluripotency and differentiation markers behaved as in *Eed*^-/-^ cells without PE deletion (Fig S5D). We conclude that counteracting H3K27me3 is not the main role of PE in *Fgf5* regulation.

### PE does not affect activation of the intergenic enhancers

Studies on the Wap-SE have suggested that individual elements can affect the activation of unrelated elements within the same cluster (Shin *et al*., 2016). We therefore performed ChIP-qPCR to test whether H3K27ac accumulation at the E1 or E2 enhancer was similarly affected by deletion of PE. We detected similar amounts of H3K27ac at the E1 and E2 enhancers in WT and ΔPE cell lines at 40 h of differentiation (Fig 5C). However, loss of E1 affected H3K27ac deposition at the E2 enhancer (Fig 5C), and we observed reduced H3K27ac levels at the E1 enhancer upon E2 deletion (although not significant, p-value=0.06). H3K27ac accumulation at control enhancers was comparable between the different cell lines (Fig S5F). We conclude that E1 and E2 are activated independently of PE, but affect each other’s activation status.

### Accumulation of Pol II at PE

Next, we tested whether loss of PE indeed reduces *Fgf5* transcription or whether it decreases mRNA stability through unknown mechanisms without affecting transcription. To analyze nascent transcription, we performed PRO-Seq (Mahat *et al*., 2016) 40 h post-differentiation, comparing WT, ΔPE, and all the PE KI cell lines (Fig 6A). Nascent transcription around the TSS as well as the first exon might be confounded by divergent transcription originating at PE and might not be suitable to compare WT and ΔPE cell lines with each other. Therefore, we quantified Spike-In normalized nascent transcript levels across the second and third exon of *Fgf5* to compare overall levels of transcription (Fig 6B).

**Figure 6:**
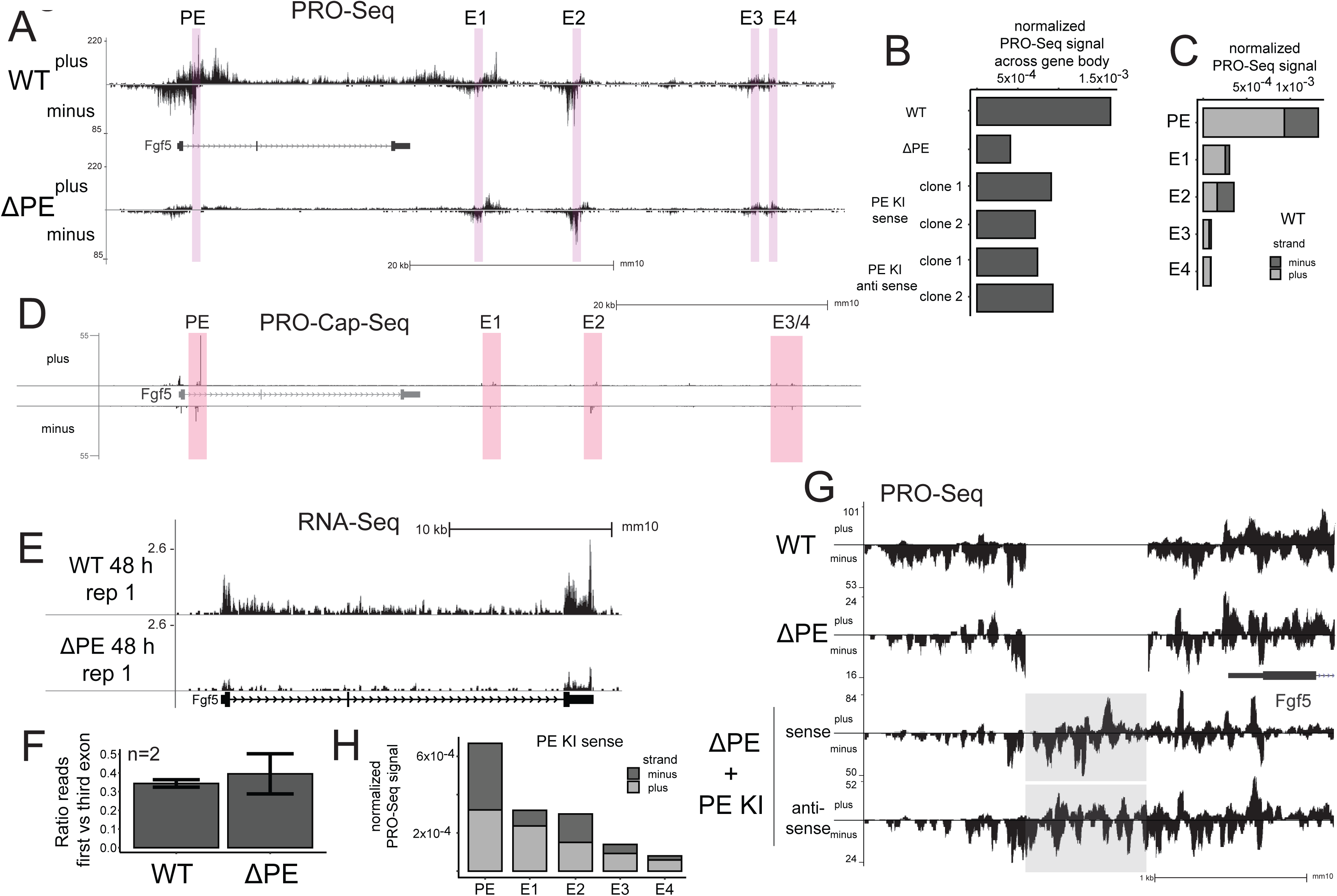
High levels of Pol II accumulate at the PE element. (A) Spike-In-normalized strand-specific PRO-Seq signal at the *Fgf5* locus in WT and ΔPE cells after 40 h of ESC to EpiLC differentiation with fixed scale bar. Enhancers are highlighted in red. (B) Quantification of Spike-In-normalized PRO-Seq signal on the plus strand between start of *Fgf5* exon two and end of *Fgf5* exon three in WT, ΔPE as well as PE KI cells after 40 h of ESC to EpiLC differentiation. (C) Quantification of Spike-In-normalized PRO-Seq signal at the *Fgf5* enhancers on plus and minus strand in WT cells after 40 h of differentiation. (D) Spike-In-normalized strand-specific PRO-Cap-Seq signal with nucleotide resolution at the *Fgf5* locus in WT cells after 40 h of ESC to EpiLC differentiation. Enhancers are highlighted in red. (E) RiboZero RNA-Seq signal normalized for sequencing depth at the *Fgf5* locus in WT and ΔPE cells after 48 h of ESC to EpiLC differentiation with fixed scale bar. For the second replicate and a representation with adjusted scale bar, see Supplements. (F) Quantification of RiboZero RNA-Seq signal in *Fgf5* exon one divided by *Fgf5* exon three in WT and ΔPE cells after 48 h of ESC to EpiLC differentiation. Mean values of n=2 biological replicates are shown. Error bars correspond to one standard deviation in each direction. (G) Spike-In-normalized strand-specific PRO-Seq signal at the *Fgf5* locus in WT, ΔPE and PE KI (one anti-sense and one sense clone, see Supplements for additional clone) cells after 40 h of ESC to EpiLC differentiation with adjusted scale bar. The knocked-in PE element is highlighted in grey. (H) Quantification of Spike-In-normalized PRO-Seq signal at the *Fgf5* enhancers on plus and minus strand in PE KI (sense) cells after 40 h of ESC to EpiLC differentiation. For similar quantifications in the remaining clones, see Supplement.

Loss of PE indeed reduced nascent transcription compared to WT. This reduction was partially rescued in the KI cell lines, albeit not to WT levels (Fig 6A, 6B, S6A). Transcription across the *Pou3f1*/*Oct6* gene was comparable between all cell lines (Fig S6B). Next, we calculated the travel ratio of Pol II in each of the WT and mutant cell lines by dividing PRO-Seq reads in the gene body by those mapping close to the TSS (Fig S6C). Even though loss of PE decreased nascent *Fgf5* transcription, it did not affect the ratio between initiating and actively transcribing Pol II. From these data, we conclude that PE indeed contributes to *Fgf5* transcription, without affecting promoter-proximal pausing.

When comparing the PRO-Seq tracks, we noticed a stronger accumulation of nascent transcript at PE compared to the promoter (Fig 6A), reminiscent of a paused polymerase peak at the enhancer. It has been previously shown that most enhancers show some Pol II transcription leading to the production of short-lived eRNAs (Kim *et al*., 2010; Schwalb *et al*., 2016). Indeed, we also observed active transcription at all enhancers analyzed in this study (Fig 6A, S6A and S6B). However, the levels of Pol II at PE were 5- to 10-fold higher compared to E1 through E4 (Fig 6C). We validated the accumulation of Pol II at PE during differentiation, using an independently derived publicly available Pol II ChIP-Seq dataset (Yang *et al*., 2019) (Fig S6E). Starting from 24 h, Pol II accumulated at PE and at the TSS, but only to a much lower degree at E1 through E4.

Next, we analyzed the origin of Pol II at the PE element. Pol II initiating at the promoter could be stalled at PE. Alternatively, Pol II could be recruited directly to PE and initiate at an alternative TSS, as has been described previously (Kowalczyk *et al*., 2012). To distinguish between these two possibilities, we performed PRO-Cap-Seq (Mahat *et al*., 2016) to enrich for capped nascent transcripts and determine the exact site of transcription initiation by sequencing them from the 5’-end. Using this technique, we found some signal at the promoter, but we also observed a very strong and distinct peak at PE (Fig 6D). The PRO-Cap-Seq signal at PE was again much stronger than the signal at E1-E4. These results suggest that PE serves as a strong transcription initiation site, thus accumulating Pol II.

We conclude that accumulation of high levels of Pol II at PE is due to initiation directly at the PE element. As PE is positioned within an intron or upstream of the promoter in case of the KI cell lines, Pol II initiating at PE might in both cell lines proceed to productive elongation and give rise to *Fgf5* mRNA. Therefore, PE might act as an alternative promoter, rather than as an enhancer that activates transcription from the endogenous promoter. However, the RiboZero RNA-Seq signal in WT cells at the PE element was much lower compared to the signal at the *Fgf5* exons (Fig 6E and S6D). Exon two showed relatively low signal, probably because of the existence of an isoform containing only exons one and three.

Pol II that initiates at PE and continues to transcribe through the entire gene would contribute to RNA-Seq reads downstream of the PE (i. e. in exon two and three), but not upstream of it in exon one. Therefore, deletion of PE and removal of this putative alternative promoter should reduce RiboZero RNA-Seq reads in the third exon more strongly than in the first exon. Similarly, nascent transcription downstream of PE should be more severely affected by PE deletion than nascent transcription upstream of PE. However, neither the ratio of RNA-Seq reads between exons one and three nor the travel ratio of PRO-Seq reads in the gene body compared to the TSS were significantly affected by deletion of PE (Fig 6F and S6C). In addition, the read coverage was similarly reduced across the entire *Fgf5* locus upon deletion of PE (Fig 6E and S6D), although we do note that the sparse coverage due to lower expression levels upon deletion of PE might exacerbate visual analysis of RNA-Seq tracks. Finally, the forward primer used for RT-qPCR analysis of *Fgf5* expression (Fig 3A and 3B) maps to the end of exon one, i. e. upstream of a potential transcript originating from PE. Therefore, the reduced expression observed upon PE deletion cannot be explained by loss of transcripts originating from PE, as those transcripts would not have been amplified by the qPCR primers. All in all, while we cannot completely rule out that some initiation at PE might give rise to a mature *Fgf5* transcript, our results indicate that PE deletion mainly affects initiation at the endogenous promoter, and that initiation at PE mostly produces short-lived transcripts, as it has been reported for eRNAs.

After identifying a strong signal of paused Pol II at PE without associated mature transcript, we wondered whether this might be the main function of PE: recruitment of Pol II at PE leading to a pool of polymerase and a higher local concentration that could be used by E1-E4 for initiation at the actual *Fgf5* promoter. Accumulation of Pol II at PE could either be an intrinsic property of the enhancer or a mere consequence of its position within an intron, where it might as well accumulate Pol II originating from the promoter. While the PRO-Cap-Seq results support the former explanation, we further tested these two scenarios by analyzing whether KI of PE 5’ of the promoter would also lead to a higher local accumulation of paused Pol II at the PE element. To account for the genetic changes in the KI cell lines, we mapped reads to custom-made bowtie indexes, in which PE had been removed from its endogenous position, and instead had been reintroduced upstream of the promoter in either sense or antisense orientation.

Indeed, in cell lines with the PE element 5’ of the promoter we found high levels of nascent transcription at PE (Fig 6G). We quantified the overall signal of nascent transcripts at PE in the KI cell lines and compared it to the extent of nascent transcripts at the intergenic enhancers E1 and E2. The overall levels of nascent transcription at E1 and E2 were slightly reduced compared to WT in all the different cell lines (Fig S6H), while transcription at the *Pou3f1/Oct6* enhancer was comparable across most cell lines (Fig S6G). However, comparisons within each cell lines showed that the strongest Pol II accumulation always occurred at PE, independent of its location within the genome (Fig 6C, 6H, S6F). The fact that accumulation of Pol II in the KI cell lines was not as strong as in WT cell lines might explain why KI of PE upstream of the promoter only partially rescued *Fgf5* expression (Fig 6B and S6A). We conclude that PE itself is recruiting higher levels of Pol II than all other enhancers within this cluster independent of its genomic location, and we hypothesize that this is important for amplification of *Fgf5* expression levels by promoting initiation at the promoter (see Discussion).

## Discussion

The study of SEs has provided conflicting results in the past. On the one hand, the individual elements within an SE have been suggested to work together in a highly cooperative fashion to activate their target genes, potentially via phase separation driven by high concentrations of TFs, co-factors and Pol II (Hnisz *et al*., 2017). Other studies suggested that each enhancer element acts independently of the others and contributes to target gene expression in an additive manner (Hay *et al*., 2016), while non-SE elements were also reported to have strong effects on target gene expression (Moorthy *et al*., 2017). To address the temporal contribution and cooperativity of individual enhancer elements to the overall expression of their target gene, we genetically dissected the *Fgf5* enhancer cluster during the differentiation of ESCs to EpiLCs. We demonstrate that the different enhancer elements at the *Fgf5* locus contribute to *Fgf5* expression at distinct time points in a super-additive manner (Bothma *et al*., 2015), and we suggest that our observations can be explained by a new mechanism of action for the PE amplifier element that involves accumulation of Pol II.

We decided to focus our study on the *Fgf5* locus due to its lack of impact on early embryonic development, as it allows a detailed analysis of enhancer deletions and their effect on target gene expression during cell fate transition without perturbing the differentiation process itself. Epigenomic mapping through ChIP-Seq analysis against p300, H3K4me1 and H3K27ac at 48 h of differentiation had previously identified five individual putative enhancer elements at the *Fgf5* locus (Buecker *et al*., 2014). While the intronic PE element seems to amplify *Fgf5* expression at all time points and its loss lead to a general shift of the *Fgf5* expression curve towards lower expression levels, the four intergenic elements are controlling the induction of *Fgf5* expression during the exit from naïve pluripotency. These intergenic elements showed different dynamics: loss of E1 lead to the earliest reduction in *Fgf5* expression compared to WT, followed by E2 and finally E3 and E4.

Interestingly, these dynamics were not reflected by the acquisition of the active enhancer mark H3K27ac. Here, E3 gained H3K27ac before E1 and E2, however, loss of E3 only affected *Fgf5* expression at a later stage compared to loss of E1 and E2. Conversely, deletion of the PE element reduced *Fgf5* expression levels before this enhancer accumulated noteworthy levels of H3K27ac. Our results raise the question of how instructive H3K27ac is for enhancer function, especially along a differentiation time course with high temporal resolution. It has recently been reported that H3K27ac is dispensable for ESC identity and enhancer activation (Zhang *et al*., 2020), however, differentiation analysis was not included in this report.

Similarly, only a subset of putative enhancer elements defined by epigenomic analysis consistently activated transcription in massively parallel reporter assays (MPRAs) (Barakat *et al*., 2018; Catarino & Stark, 2018). All in all, our results indicate that deposition of H3K27ac does not directly report on the actual timing of the activity of the specific enhancer. It can occur either earlier (as seen for E3) or later (as seen for PE). It is tempting to speculate that the E3 enhancer might be actively repressed early in differentiation and that it can only contribute to *Fgf5* expression upon removal of this repressor. Alternatively, the genomic distance rather than the exact timing of H3K27ac accumulation might determine when an enhancer contributes to *Fgf5* expression, as deletion of those enhancers that are closest to the promoter (PE, E1) also showed the earliest effect and *vice versa*. While enhancer activity is generally believed to be independent of genomic distance and large distances can be overcome by enhancer-promoter loops (Furlong & Levine, 2018), recent studies suggest that enhancer-promoter distance can indeed have an effect on expression levels (Carleton *et al*., 2017; Scholes *et al*., 2019). Future studies will show whether the distance between enhancer and promoter can also affect the timing of enhancer activity in a developmental setup. Importantly, the discrepancy between the timing of H3K27ac accumulation at an enhancer element and reduced target gene expression upon its deletion could only be detected by following activation of an enhancer cluster during a cell fate transition with high temporal resolution.

PE and the outside enhancers act in a super-additive manner, as expression levels of a PE only cell line and a ΔPE cell line did not add up to WT levels. Previous studies in Drosophila have suggested that multiple weak enhancers could act simultaneously at a promoter to achieve higher or super-additive transcription initiation rates compared to individual enhancers (Bothma *et al*., 2015; Carleton *et al*., 2017). To exclude that the observed super-additive effect between PE and the outside enhancers is caused by disruption of the intron and/or lower RNA stability upon deletion of PE, we transplanted this element upstream of the promoter, where it restored expression almost to WT levels.

It has been previously suggested that bidirectional transcription from intronic enhancers could negatively regulate expression of the host gene through transcriptional interference (Cinghu *et al*., 2017). When placing PE outside of the intron and upstream of the promoter, this attenuating effect should be relieved and the resulting expression levels should be higher than in a WT cell line. However, KI of PE upstream of the promoter only partially restored WT expression levels. Whether this means that transcriptional interference does not play a role at the *Fgf5* locus or whether additional surrounding sequences within the intron provide a more active environment for the PE element remains to be determined. Nonetheless, the fact that PE restored *Fgf5* expression from an exogenous location along with the observation that nascent transcription levels were reduced upon deletion of PE, confirms that PE indeed exerts its function of controlling *Fgf5* expression by regulating the process of transcription.

How can the super-additive behavior between PE and the outside enhancers be explained then? The individual elements of the *Fgf5* enhancer cluster showed very low enhancer activity in classical luciferase assays, even when combined with the endogenous promoter. Hence, enhancer-promoter incompatibilities as described between developmental enhancers and housekeeping promoters (Zabidi *et al*., 2015) do not explain these low activities. While we do note that the luciferase assays in differentiating cells suffer from high variability between biological replicates, we were able to show significant enhancer activity for the *Pou3f1/Oct6* enhancer, but not for any of the *Fgf5* enhancers. This discrepancy between the strong reduction of *Fgf5* expression upon deletion of the enhancers at the endogenous locus and their low activity in luciferase assays was especially evident for PE. While discrepancies between enhancer activity in luciferase assays and reduction of target gene expression upon deletion at the endogenous locus have been reported previously (Hnisz *et al*., 2015), a detailed mechanistic explanation for this phenomenon is still missing. Here, we suggest that PE might activate transcription at the endogenous locus via a novel mechanism that is not reflected in luciferase enhancer assays.

This novel mechanism might hinge on the enrichment of higher levels of Pol II at PE compared to E1 through E4. This accumulation of Pol II at PE could be the result of binding of a specific combination of TFs and co-activators that remain to be identified. Alternatively, presence of an enhancer with open chromatin close to the promoter – as it is the case at both the endogenous location and in the KI cell lines – might be sufficient to result in Pol II accumulation, similarly but to lower levels than what has been described in the case of Herpes Simplex Virus infection (McSwiggen *et al*., 2019). Polymerase undergoing termination or being released from DNA after promoter-proximal pausing (Steurer *et al*., 2018) might therefore be trapped at the *Fgf5* locus by PE and thus undergo several, rather than a single round of transcription (J. Li *et al*., 2019), before being released from the locus.

Accumulation of Pol II at PE might enable it to amplify expression at the *Fgf5* locus in combination with the outside enhancers. In this model, Pol II initiates at the PE element and pauses close to the initiation site but does not proceed to active elongation. According to previous studies, paused Pol II is not a stable complex bound to DNA for long periods of time, but rather quickly dissembled (Erickson *et al*., 2018; Krebs *et al*., 2017; Steurer *et al*., 2018). This removal of paused Pol II from DNA might be actively regulated by the Integrator complex (Elrod *et al*., 2019; Tatomer *et al*., 2019). In our model, paused Pol II that is quickly released from the PE element accumulates in the vicinity of the *Fgf5* promoter. This pool of accumulated Pol II can subsequently be recruited to the promoter for initiation and production of an mRNA. PE thus amplifies the contribution of the other regulatory elements at the locus - in this case the Fgf5 promoter as well as E1-E4 - in a super-additive fashion by increasing the local concentration of Pol II.

In conclusion, we suggest that PE does not function as a canonical enhancer, but rather as an “amplifier” of overall levels of transcription at the *Fgf5* locus. Detection of this amplifier element was only made possible through carefully dissecting the contribution of individual putative enhancer elements to their target gene expression along a differentiation time course. We envision that similar studies at individual loci will identify additional amplifier elements and resolve whether all epigenomically identical enhancers activate transcription by the same mechanism.

## Acknowledgments

We would like to thank all members of the Buecker lab for discussions and feedback throughout the project, Ursula Schöberl for technical help with establishing PRO-Seq and PRO-Cap-Seq, Alexander Stark for critical feedback and discussions on the manuscript, the BioOptics facility at Max Perutz Labs as well as the NGS facility at VBCF. This work was supported by the FWF (P 30599 to C.B.) and Uni:Docs fellowships from the University Vienna to H.T. and M.R.

## Methods

### ESC maintenance

Mouse ESCs were cultured in base medium - HyClone^TM^ DMEM/F12 medium without HEPES (GE Healthcare) with 4 mg/mL AlbuMAX^TM^ Lipid-Rich Bovine Serum Albumin (Gibco^TM^), 1x serum-free B-27^TM^ Supplement (Gibco^TM^), 1x N2 supplement (homemade, components purchased from Sigma-Aldrich and R&D Systems), 1x MEM NEAA (Gibco^TM^), 50 U/mL Penicillin-Streptomycin (Gibco^TM^), 1 mM Sodium Pyruvate (Gibco^TM^) and 1x 2-Mercaptoethanol (Gibco^TM^) - supplied with 3.3 μM CHIR-99021 (Selleckchem), 0.8 μM PD0325901 (Selleckchem) and 10 ng/mL hLIF (provided by the VBCF Protein Technologies Facility, www.vbcf.ac.at) (from here on referred to as 2i/LIF medium) on CELLSTAR® 6-well plates (Greiner Bio-One) coated first with Poly-L-ornithine hydrobromide (6 μg/mL in 1xPBS, 1 h at 37 °C, Sigma-Aldrich) and then with Laminin from Engelbreth-Holm-Swarm murine sarcome basement membrane (1.2 μg/mL in 1xPBS, 1 h at 37 °C, Sigma-Aldrich). They were passaged every two to three days in an appropriate ratio. Therefore, 250 μL of 1x Trypsin-EDTA solution (Sigma-Aldrich, T3924) were used and trypsination was stopped with 2i/LIF medium containing 10% Fetal Bovine Serum (Sigma-Aldrich, F7524).

### Generation of KO&KI cell lines

For deleting a given enhancer, two gRNAs targeting the left and right boundary of their respective p300 ChIP-Seq peak (data from Buecker *et al*., 2014) were designed with CRISPRscan (Moreno-Mateos *et al*., 2015). Forward and reverse DNA oligonucleotides - containing the gRNA-Sequence as well as the overhangs required for cloning - were ordered from Microsynth AG, annealed and cloned into BbsI-digested (NEB) pX330-U6-Chimeric_BB_CBh_hSpCas9 plasmid (Cong *et al*., 2013). The resulting plasmids expressed the gRNA from a U6 promoter and the Cas9 protein from the CBh promoter.

200,000 mouse ESCs were seeded in one well of a 6-well plate and on the following day transfected with 950 ng of each gRNA-containing plasmid as well as 100 ng of plasmid expressing a fluorescent marker. Therefore, Lipofectamine® 2000 Transfection Reagent (Thermo Fisher Scientific) was used. The three plasmids were diluted in 100 μL of DMEM/F12 medium, and 12 μL of transfection reagent were diluted in 100 μL of DMEM/F12 medium. After 5 minutes (min) of incubation at room temperature, the diluted plasmids were added drop wise to the DMEM/F12-transfection reagent mixture. After another 30 min incubation at room temperature, this transfection mix was added drop wise to the cells. 6-8 h after adding the transfection mix, the medium was removed and fresh 2i/LIF medium added to the cells.

Two days after transfection, a single fluorescent cell was sorted per well of a fibronectin-coated (10 μg/mL Human Plasma Fibronectin Purified Protein (Sigma Aldrich) in 1x PBS, 1 h at 37 °C) 96-well plate. As sub-stoichiometric amounts of plasmid expressing the fluorescent marker had been transfected, cells carrying this fluorescent marker are highly likely to also carry the gRNA-expressing plasmids. Deletion of the respective enhancer was confirmed by PCR with primers mapping outside of the sites recognized by the two gRNAs, thus giving rise to shortened PCR product in case of successful deletion.

For generating enhancer KIs, an enhancer sequence similar in size to what had been deleted in the respective KO cell line was amplified by PCR either in sense or in antisense orientation, and cloned into an AgeI-HF®- and XbaI-digested (both NEB) pGemT-plasmid containing a puro-delta TK selection cassette surrounded by loxP sites. Left and right homology arms targeting the desired KI site in the genome were designed to be 800-900 bp long, and to be separated by roughly 30 bp. They were amplified by PCR and inserted upstream of the enhancer and downstream of the second loxP site by Gibson assembly, respectively. After assembly of this plasmid – containing left and right homology arm, the enhancer as well as the loxP-flanked selection cassette – it was linearized by restriction digestion.

A single gRNA targeting the genomic sequence between left and right homology arm was designed and cloned into the pX330-U6-Chimeric_BB_CBh_hSpCas9 plasmid as described above.

200,000 mouse ES cells were seeded in a 6-well and on the following day transfected with 400 ng of linearized plasmid as well as 400 ng of gRNA-containing plasmid, as described above. One day after the transfection, cells were passaged and transferred onto a 10 cm dish. Within 48 h of the transfection, positive integration events were selected for with puromycin (2 μg/mL, InvivoGen). Single colonies were picked into fibronectin-coated (10 μg/mL) 96-well plates after one week of selection, and correct integration was validated by PCR.

Colonies with correct integration and intact homology arms were expanded and transfected with plasmid expressing Cre-recombinase to remove the selection cassette as described above (200,000 cells, 1 μg of Cre-recombinase expressing plasmid, 5 μL of transfection reagent). Cells were passaged and seeded at low density on the day after transfection. Selection with ganciclovir (500 ng/mL, Invivogen) for successful removal of the selection cassette was started within 48 h of the transfection. After one week of selection, single colonies were picked and removal of the selection cassette was confirmed by PCR (PE KI validation 1 primers). In addition to this, KI of the enhancer and intactness of the homology arms was confirmed by PCR using primers mapping outside of the left and right homology arms respectively (PE KI validation 2 primers), and subsequent Sanger sequencing of the PCR product.

### Differentiation and RT&qPCR analysis

For differentiation and subsequent RT-qPCR analysis, 100,000 cells per cell line and time point were seeded in 2i/LIF medium on fibronectin-coated (5 μg/mL) 12-well plates. On the following day, the medium was removed and cells were washed twice with 1 mL of 1x PBS. 1 mL of base medium supplied with 12 μg/mL Recombinant Human FGF-basic (PEPROTECH) and KnockOut^TM^ Serum Replacement (1:100, Gibco^TM^) (from here on referred to as FK medium) was added to start differentiation; for the 0 h time point, 1 mL of fresh 2i/LIF medium was added.

After 12, 24, 36 and 48 h of differentiation, cells were lysed in 500 μL of pepGOLD TriFast^TM^ reagent (Peqlab) and stored at −80 °C until ensuing RNA extraction. For the 0 h time point, samples were collected 48 h after adding fresh 2i/LIF medium. RNA was extracted by phenol-chloroform extraction, precipitated with Isopropanol and washed with 75% ethanol according to the pepGOLD TriFast^TM^ extraction protocol. RNA was re-suspended in 15 μL of RNase free water and subsequently quantified. 800 ng of RNA were used for reverse transcription with the SensiFAST^TM^ cDNA Synthesis kit (Bioline) according to the standard protocol.

For subsequent qPCR analysis with the SensiFAST^TM^ SYBR® No-ROX kit (Bioline), 0.5 μL of resulting cDNA were used per 10 μL reaction along with 125 nM of forward and reverse primer. qPCR primers were designed with Primer3 (Koressaar & Remm, 2007). qPCR reactions were performed in technical triplicates following the recommended 2-step cycling qPCR programme.

For each primer, time point and cell line, mean Cq values were calculated based on the technical triplicates. ΔCq values were calculated by subtracting the mean Cq value of the primer of interest from the mean Cq value of the Rpl13a primer, and normalized expression values were calculated by 2^ΔCq^. For each cell line, biological replicates were performed independently (i. e. cell lines were seeded and differentiated on different days) and for each experiment a WT cell line was included. Mean normalized expression values were calculated and are depicted in line graphs (see Figures).

For quantitative analysis and statistical testing, expression values of each cell line and time point were normalized to the expression values of the WT cell line from the same experiment at the corresponding time point. The resulting values were then averaged across the biological replicates and are depicted in bar graphs (see Figures).

In addition to this, for *Fgf5* expression values a one-sided Welch Two sample t-test was performed on these WT-normalized values to assess whether they are significantly lower (or in rare cases higher) than 1 (as all values are normalized to WT, a value of 1 corresponds to WT expression levels). For control genes, a two-sided Welch Two sample t-test was performed on the WT-normalized values to assess whether they are significantly different from 1. In both cases, p-values lower than 0.05 were regarded as statistically significant.

Statistical analysis was performed in R version 3.6.3 (R Core Team, 2013) and graphs were generated with the ggplot2-3.3.0 package (Wickham, 2016).

### RiboZero RNA-Seq

Cells were differentiated and RNA extracted from two biological replicates as described above. RNA-Seq libraries depleted for ribosomal RNA were generated and sequenced at the VBCF NGS Unit (www.viennabiocenter.org/facilities).

Libraries were sequenced to a depth of 23-27 million reads (single-end, 50 bp). Adapters were removed with the adapter auto-detection function of Trim Galore Version 0.5.0 (https://github.com/FelixKrueger/TrimGalore) and reads were aligned to the mm10 assembly of the mouse genome (downloaded from https://www.encodeproject.org/data-standards/reference-sequences) using the splice-sensitive STAR_2.5.3a aligner STAR (Dobin *et al*., 2013). SAMtools 1.5 (H. Li *et al*., 2009) was used to sort and index the resulting bam files, as well as for extracting uniquely mapping reads.

Reads mapping to the exon of each gene were counted with the featureCounts function of the Rsubread package (version 1.5.3) (Liao *et al*., 2019). Differentially expressed genes (log2fold change of bigger than 1 or lower than −1; adjusted p-value of 0.05 or lower) were identified with the DESeq2 package 1.26.0 (Love *et al*., 2014).

### SMART-Seq2 single-cell RNA-Seq

100,000 WT cells were seeded in 2i/LIF medium on fibronectin-coated (5 μg/mL) 12-well plates. Differentiation was started at staggered time points to allow for sample collection in parallel at the same time (earliest 4 h post seeding). Therefore, cells were washed with 1 mL of 1x PBS, and FK medium was added. Single cells were FACS-sorted directly into 96-well plates containing smartseq2 lysis buffer (48 cells/condition) based on forward/sideward scatter index sorting. Samples were stored at –80 °C until library preparation. To control for successful sorting, qPCRs against Rpl13a and Oct4 were performed after cDNA synthesis. Only wells, where amplification occurred, were selected for further library preparation (24 cells per condition). Samples were multiplexed and sequenced on two lanes of a HiSeq 3000/4000 machine (single-end, 50 bp).

Raw unaligned bam files were converted to fastq files with SAMtools 1.5 (H. Li *et al*., 2009). Reads were aligned to Mus_musculus.GRCm38.90 with the splice-sensitive STAR_2.5.3a aligner (Dobin *et al*., 2013) and aligned reads were counted with the featureCounts function of the Rsubread package (version 1.5.3) (Liao *et al*., 2019). After generating the counttable, data was analysed with the Bioconductor SingleCellExperiment workflow (Lun & Risso, 2019) and scater (McCarthy *et al*., 2017). Cells were filtered based on library size and mitochondrial content.

### Luciferase assays

For luciferase assays, we used a pGL3-plasmid with the Firefly luciferase coding sequence followed by a poly-adenylation signal under the control of a SV40 promoter. Enhancers fragments were defined based on p300 and OCT4 as well as OTX2 ChIP-Seq data (Buecker *et al*., 2014), amplified by PCR and inserted downstream of the poly-adenylation signal by Gibson assembly. For assays with the endogenous promoter, the SV40 promoter was removed from the luciferase-enhancer plasmids by restriction digestion with BglII and HindIII-HF (both NEB). The *Fgf5* promoter region - encompassing the 300 bp region containing most of transcription initiation events in PRO-Cap-Seq data at the 5’ UTR of the gene plus 100 bp of flanking nucleotides on each side - was amplified by PCR and inserted in place of the SV40 promoter by Gibson Assembly. In cases, where either restriction enzyme motif was also present in the respective enhancer, we first substituted the promoter in the luciferase plasmid without enhancer, and then added the enhancers from scratch.

To control for differences in transfection efficiency, we co-transfected a plasmid constitutively expressing Renilla luciferase. As Firefly and Renilla luciferase have different substrate specificity and different optimal reaction conditions, luciferase activity of the two enzymes can be measured independently.

For luciferase assays, 5,000 cells were seeded per well of a fibronectin-coated (10 μg/mL) 96-well plate. On the following day, cells were transfected with 20 μL of transfection mix containing 120 ng of enhancer-luciferase plasmid, 4 ng of Renilla control plasmid and 0.62 μL of Lipofectamine® 2000 Transfection Reagent. Luciferase assays were performed in technical triplicates, i. e. for each plasmid and time point 3 wells of cells were transfected. In addition to this, 3 wells of untransfected cells and 3 wells transfected with no-enhancer control (luciferase plasmid containing the respective promoter, but no additional enhancer) were included in every experiment for background subtraction and normalization.

5-7 h after transfection, the medium was removed and cells were washed twice with 150 μL 1x PBS. 175 μL FK medium were added to start differentiation. 24 or 40 h after starting the differentiation, luciferase activity was measured using the Dual-Glo® Luciferase Assay System (Promega). Therefore, the medium was removed and 40 μL of fresh FK medium were added. Cells were incubated at room temperature for 30 min and lysed by addition of 40 μL of Dual-Glo® Reagent. After 10 min incubation at room temperature, Firefly luminescence - resulting from expression of the enhancer-luciferase plasmid - was measured. 40 μL of Dual-Glo® Stop&Glo® Reagent were added and after 10 min incubation Renilla luminescence - resulting from expression of the Renilla control plasmid - was measured.

To estimate the background for each measurement, the average value of the three untransfected wells was calculated for both the Firefly and the Renilla measurement. These background values were subtracted from the Firefly and Renilla measurements of the transfected cells respectively. To normalize for transfection efficiency, for each well the Firefly measurement was normalized to the Renilla measurement (as identical amounts of Renilla plasmid were transfected for every well, differences in Renilla signal reflect different transfection efficiencies). The resulting values were averaged across the technical triplicates. Subsequently, they were normalized to the no-enhancer control, in which luciferase expression was driven by the same promoter in the absence of any additional enhancer. As insertion of enhancers increases the molecular weight of the plasmids, identical masses of plasmid (in our case 120 ng) contain different numbers of plasmid molecules, i e. for bigger plasmids less molecules had been transfected. To account for this, we normalized the size of each enhancer-luciferase plasmid to the no-enhancer control, and multiplied the no-enhancer normalized values of luciferase activity with this factor.

For each plasmid, biological replicates were performed independently (i. e. cells were seeded, transfected and differentiated on different days). The values normalized for no-enhancer control and plasmid-size were averaged across the biological replicates, and they were also used to assess statistical significance. Therefore, a two-sided Welch Two sample t-test was performed to test whether these values are significantly different from 1 (a value of 1 corresponds to the luciferase activity driven by the promoter only in the absence of any enhancer). p-values lower than 0.05 were regarded as statistically significant.

## H3K27ac-ChIP

### Differentiation of cells and collection of ChIP pellets

For the H3K27ac ChIP-Seq time course, 3,000,000 cells were seeded per fibronectin-coated (5 μg/mL) 15 cm dish. On the following day, the medium was removed and cells were washed twice with 15 mL of 1x PBS. 20 mL of FK medium were added to start differentiation; for the 0 h time point, 20 mL of fresh 2i/LIF medium were added. Samples were collected after 12, 18, 24, 30, 36, 43 and 48 h of differentiation. For the 0 h time point, samples were collected 48 h after adding fresh 2i/LIF medium.

In case of all other ChIPs, cells were passaged and resulting cell pellets were washed twice with 10 mL of base medium. Cells were resuspended in base medium and 3,000,000 cells per fibronectin-coated (5 μg/mL) 15 cm dish were directly seeded in either FK medium (for differentiated samples) or 2i/LIF medium (for undifferentiated samples). Samples were collected 40 h after plating.

Therefore, the medium was removed and 10 mL of 1x PBS were added. Formaldehyde was added to a final concentration of 1% to cross-link proteins to DNA. After 10 min incubation at room temperature, glycine was added to a final concentration of 0.125 M to quench the formaldehyde. After another 10 min incubation at room temperature, the PBS/formaldehyde/glycine mixture was removed and cells were washed twice with 10 mL of cold 1x PBS. 10 mL of cold 1x PBS with 0.01% of Triton X-100 were added and cells were collected with a cell scraper. After centrifugation at 4 °C and 500 g for 5 min, the supernatant was discarded, cell pellets were flash frozen in liquid nitrogen and stored at −80 °C. As for the H3K27ac time course the size of the cell pellets varied between the different time points, pellets from multiple plates were pooled and the size of the cell pellets manually adjusted to the size of the pellet for the 48 h time point. For all other ChIPs, one pellet was collected per 15 cm dish.

### ChIP

Pellets were thawed on ice for 30 min, resuspended in 5 mL cold LB1 buffer (1 M Hepes-KOH pH 7.5, 5 M NaCl, 0.5M EDTA, 50% gylcerol, 10 %NP-40, 10% Triton X-100, 1 mM PMSF, 1x cOmplete^TM^ Protease Inhibitor Cocktail (Roche)) and rotated for 10 min at 4 °C. After centrifugation for 5 min at 1350 g and 4 °C, the supernatant was removed, and the pellet was resuspended in 5 mL cold LB2 buffer (1 M Tris-Hcl pH 8.0, 5 M NaCl, 0.5 M EDTA, 0.5 M EGTA, 1mM PMSF, 1x cOmplete^TM^ Protease Inhibitor Cocktail) as well as rotated for 10 min at room temperature. After another centrifugation for 5 min at 1350 g and 4 °C, the supernatant was removed once more and the pellet resuspended in 1.5 mL cold LB3 buffer (1 M Tris-HCl pH 8.0, 5 M NaCl, 0.5 M EDTA, 0.5 M EGTA, 10% sodium deoxycholate, 20% N-lauroylsarcosine, 1 mM PMSF, 1x cOmplete^TM^ Protease Inhibitor Cocktail). Samples were sonicated in 15 mL Bioruptor® Pico Tubes (diagenode) with 200 μL of sonication beads (diagenode) in a Bioruptor® Pico sonication device (diagenode) for 14 cycles with 30 s on and 45 s off, and transferred to a fresh 1.5 mL reaction tube. After centrifugation for 10 min at 16000g and 4 °C, the supernatant was transferred to a new tube and 150 μL of 10% Triton X-100 were added.

500 μL of chromatin and 5 μg of antibody (Histone H3K27ac antibody (pAb),Active Motif (39133)) were used per cell line and time point. After adding the antibody, samples were rotated at 4 °C overnight to bind the antibody to the chromatin. 50 μL of sonicated chromatin were used as Input samples and stored at −20 °C.

On the following day, 100 μL of Protein G Dynabeads (Dynabeads^TM^ Protein G for Immunoprecipitation, Thermo Fisher Scientific) were aliquoted per ChIP-sample and washed three times with 1 mL of cold block solution (0.5% BSA in 1x PBS), to block unspecific binding to the beads. Chromatin was added to the beads, and samples were rotated at 4 °C for 4 h to allow for binding of antibody-bound chromatin to the beads.

Bound beads were washed five times with 1 mL of cold RIPA buffer (1 M Hepes-KOH pH 7.5, 5 M LiCl, 0.5 M EDTA, 10% NP-40, 10% sodium deoxycholate) and one time with cold 1x TE + 50 mM NaCl. After centrifugation for 3 min at 950 g and 4 °C, all remaining supernatant was removed and 210 μL of elution buffer (1 M Tris-Hcl pH 8.0, 0.5 M EDTA, 10% SDS) were added. Samples were incubated at 65 °C for 15 min and briefly mixed every few minutes. After centrifugation for 1 min at 16000 g and room temperature, 200 μL of supernatant containing the eluted chromatin were transferred to a fresh tube. Input samples were thawed and 3 volumes of elution buffer were added. After brief mixing, both ChIP and Input samples were incubated at 65 °C overnight to reverse crosslinks.

On the following day, samples were diluted with 1 volume of TE buffer and RNase A (Roche) was added to a final concentration of 0.2 mg/mL. After incubation for 2 h at 37 °C, CaCl_2_-Tris HCl pH 8.0 was added to a final CaCl_2_-concentration of 5.25 mM and Proteinase K (Sigma-Aldrich) was added to a final concentration of 0.2 mg/mL. Samples were incubated at 55 °C for 30 min and transferred to Phase Lock Gel^TM^ tubes (Quantabio). To extract DNA, one volume of Phenol-Chloroform-Isoamyl alcohol (25:24:1) was added and samples were mixed by inverting. After centrifugation at 16000 g and room temperature for 5 min, another volume of Phenol-Chloroform-Isoamyl alcohol was added and samples were mixed as well as centrifuged once more for 5 min at 16000 g and room temperature. The supernatant was transferred to a fresh 1.5 mL reaction tube, and 2 volumes of cold 96% ethanol as well as 1/10th volume of 3 M NaOAc and 1.5 μL of 20 mg/mL glycogen were added.

Samples were incubated at −20 °C overnight to precipitate DNA, and then centrifuged at 16000 g and 4 °C for 30 min. The supernatant was removed and 0.5 mL of cold 70% ethanol were added to wash the pellet. After brief mixing, samples were centrifuged for 15 min at 16000g and 4 °C. All supernatant was carefully removed. The pellet was air dried for 5 min at room temperature and resuspended in 50 μL of PCR-grade water (Sigma-Aldrich).

### ChIP-qPCR

Inputs were diluted 1:10 with PCR-grade water. 0.5 μL of resulting DNA (undiluted for ChIPs, diluted for Inputs) were used per 10 μL reaction with the SensiFAST^TM^ SYBR® No-ROX kit (Bioline), along with 125 nM of forward and reverse primer. qPCR reactions were performed in technical triplicates following the recommended 2-step cycling qPCR programme. qPCR primers were designed with Primer3 (Koressaar & Remm, 2007). Primers for K27ac ChIP-qPCR were designed to target the flanking regions of the p300 peak at the respective enhancer.

For each primer and cell line, mean Cq values were calculated based on the technical triplicates. ΔCq values were calculated by subtracting the mean Cq value of the respective primer with the ChIP sample from the mean Cq value of that primer with the Input sample. As 10-fold less material was used for Input samples compared to ChIP samples, and as the Input samples were diluted 10-fold before performing the qPCR, the amount of Input material per qPCR is 100-fold reduced compared to the ChIP. Therefore, Percentage of Input enrichment was calculated by 2^ΔCq^/100.

To account for differences in ChIP efficiency, we normalized these percentage of Input values to the percentage of Input values of two negative control regions, that are known not to have any active chromatin marks in ESCs or upon differentiation based on previous ChIP-Seq experiments (Buecker *et al*., 2014).

For each cell line, biological replicates were performed independently (i. e. cell lines were seeded and differentiated on different days). Percentage of Input values normalized to the negative control regions were averaged across these biological replicates and are depicted in the bar graphs. A one-sided Welch Two sample t-test was performed to test whether these values are significantly higher or lower compared to WT. p-values lower than 0.05 were regarded as statistically significant.

### ChIP-Seq

ChIP and Input samples were quantified with a Fluorescence NanoDrop. DNA libraries were then generated on ice with the sparQ DNA Library Prep Kit (Quantabio) following the standard protocol with some modifications that are described below. Different adapters were used for each sample to allow for multiplexing samples and including them in the same sequencing run.

To avoid over-amplification of libraries, we followed a special protocol for the PCR amplification. PCR reactions were prepared as suggested in the standard protocol. However, after 5 cycles of amplification the PCR reactions were stopped and stored on ice. To estimate how many additional cycles of PCR were required for optimal library amplification, 5 μL of each library were used to prepare an additional 15 μL PCR reaction for each library that contained 0.1x SYBR® Green I nucleic acid gel stain (Sigma-Aldrich) and was run in a qPCR machine for an additional 40 cycles following the exact same protocol. Based on the relative fluorescent units measured by the qPCR, a threshold was determined for each library at 25% of saturation level, at which fluorescence did not increase with additional PCR cycles any more. We then estimated at which cycle this threshold concentration had been reached during the qPCR, and resumed PCR amplification of the original libraries for this exact number of cycles. For most libraries we performed a total of 5-8 cycles of PCR amplification.

After PCR amplification, we continued following the standard protocol, but included an additional purification step with AMPure XP beads (1.8 x, Beckman Coulter) to remove adapters and primers that remained in the supernatant, whereas the libraries bound to the beads and were eluted after removing the supernatant.

The size distribution of the libraries was analyzed with the Agilent High Sensitivity DNA kit. If necessary, additional purification with AMPure XP beads was performed to remove primers and adapters (purification with 1x AMPure XP beads; the supernatant was discarded and the DNA bound to the beads subsequently eluted) or to exclude DNA fragments of more than 1 kb (purification with 0.54 x AMPure XP beads; the high molecular weight fragments bound to the beads and were discarded, while the library enriched for smaller DNA fragments remained in the supernatant).

Libraries were quantified with the PerfeCTa® NGS Quantification kit (Quantabio) and similar amounts of each library were pooled based on this quantification for next-generation sequencing. Sequencing was performed at the VBCF NGS Unit.

Libraries were sequenced to a depth of 8-18 million reads (single-end, 50 bp). Reads with identical sequence, that are likely to be PCR duplicates, were removed with the Clumpify tool from BBTools version 37.20 (https://github.com/BioInfoTools/BBMap/blob/master/sh/clumpify.sh). Adapters were removed with the adapter auto-detection function of Trim Galore Version 0.5.0 (https://github.com/FelixKrueger/TrimGalore); in addition to this, the first two nucleotides after the adapter were also removed, as those had been artificially inserted by A-tailing during the library preparation.

Reads were aligned to the mm10 assembly of the mouse genome (downloaded from https://www.encodeproject.org/data-standards/reference-sequences) with Bowtie 2 Version 2.3.4.3 (Langmead & Salzberg, 2012). SAMtools 1.5 (H. Li *et al*., 2009) was used to convert the resulting sam files to bam files, to sort and index the bam files, as well as for extracting uniquely mapping reads. For visualization, bam files containing uniquely mapping reads were converted into bedgraph files with bedtools version 2.28.0 (Quinlan & Hall, 2010), while normalizing for sequencing depth. Bedgraph files were then converted to bigWig files using the bedGraphToBigWig (https://github.com/sccallahan/bedGraph2bigWig) tool. BigWig files were visualized with the UCSC genome browser (Kent *et al*., 2002).

### smRNA FISH

For smRNA FISH, 1,000 cells per cell line and time point were seeded in 2i/LIF medium on a fibronectin-coated (10 μg/mL) Corning^TM^ 96-well high content microplate for imaging. On the following day, cells were washed with 1x PBS, and FK medium was added to start differentiation. For undifferentiated samples, fresh 2i/LIF medium was added instead.

After 36 h of differentiation, cells were fixed with 4% formaldehyde for 30 min and subsequently washed three times with 1x PBS. FISH was performed using the QuantiGene® ViewRNA ISH Cell Assay kit. Therefore, fixed cells were treated with Detergent Solution QC for 5 min at room temperature, and then washed twice with 1x PBS. Probe sets (Fgf5 – Type 1, Tbx3 – Type 4, Otx2 – Type 6) were diluted 1:100 in pre-warmed Probe Set Diluent QF (40°C) and added to the cells. After incubation for 3 h at 40 °C, cells were washed three times with wash buffer. During each washing step, cells were incubated with the wash buffer for 2 min before removing it. PreAmplifier Mix was diluted 1:25 in pre-warmed Amplifier Diluent QF and added to the cells. Samples were incubated for 30 min at 40 °C. After washing cells three times in wash buffer – again including the 2 min incubation before removing the buffer – Amplifier Mix diluted 1:25 in pre-warmed Amplifier Diluent QF was added. Samples were incubated for 30 min at 40 °C and washed with wash buffer as described above. Label Probe Mix was diluted 1:25 in pre-warmed Label Probe Diluent QF and added to the cells. After incubation for 30 min at 40 °C in the dark, cells were washed again with a 2 min incubation for the first two wash steps and a 10 min incubation for the third. DAPI (1x in 1x PBS, Sigma Aldrich) was added to the cells and they were incubated for 2 min at room temperature, washed twice with 1x PBS and then stored in 1x PBS at 4°C until image acquisition.

For each sample, 5 to 10 pictures were acquired with a 63x oil immersion objective (Plan-Apochromat 63x/1.40 Oil DIC M27) and a 10x magnification lens. Each picture was composed of 4 dyes (DAPI - nucleus, GFP – Type 4 – TBX3, Cy3 – Type 1 – FGF5, Cy5 – Type 6 – OTX2) with a depth of 16-bit for each dye. Furthermore, each picture was taken as a Z-series through the cell body using a Zeiss LSM700 microscope.

Images were converted from czi files to tiff images with Fiji (V2.0.0-rc-65/1.52a) (Schindelin *et al*., 2012). Therefore, each czi file was split into 4 images – one for each channel (DAPI, GFP, Cy3, Cy5) – and a Z-projection was performed on each of them. The resulting files were then further processed with CellProfiler (V3.0.0) (McQuin *et al*., 2018). To estimate the number of transcripts per cell, a cellular area was defined for each cell based on the area of the nucleus as seen in the DAPI channel plus a pre-defined radius.

## PRO- and PRO-Cap-Seq

For both PRO-Seq and PRO-Cap-Seq, nuclei were isolated and nuclear run-on was performed in the exact same way (see below).

### Nuclei isolation

Cells were passaged and resulting cell pellets were washed with 12 mL of base medium. Cells were resuspended in base medium and 3,000,000 cells per fibronectin-coated (5 μg/mL) 15 cm dish were directly seeded in either FK medium (for differentiated samples) or 2i/LIF medium (for undifferentiated samples). Two plates were prepared for each cell line and condition. Samples were collected 40 h after plating.

Therefore, cells were passaged normally by adding trypsin-EDTA and stopping trypsination after incubation at 37 °C by adding base medium containing 10% serum. Resulting cell suspensions were centrifuged at 300 g and 4 °C. After removing the supernatant, cells were washed with 7.5 mL of cold 1x PBS and samples from the two plates containing identical cell line and condition were pooled. Cells were centrifuged at 300 g and 4 °C for 5 min. The supernatant was removed and cells resuspended in 1 mL of cold IA buffer (0.16 M sucrose, 3 mM CaCl2, 2 mM magnesium acetate, 0.1 mM EDTA, 10 mM TRIS-HCl pH 8.0, 0.5% NP-40; this buffer was filter-sterilized and 1 mM DTT was added directly before use). After incubation on ice for 3 min, samples were centrifuged at 700 g and 4 °C for 5 min. The supernatant was removed, samples were resuspended in 0.5 mL of cold IA buffer and incubated on ice for another 3 min. After centrifugation at 700 g and 4 °C for 5 min, the supernatant was removed, resulting nuclei were resuspended in 100 μL of cold NRB buffer (50 mM TRIS-HCl pH 8.0, 40% glycerol, 5 mM MgCl2 and 1.1 mM EDTA; this buffer was filter-sterilized) and transferred to a fresh, RNase-free 1.5 mL reaction tube. Nuclei were stained with Trypan Blue Solution (Thermo Fisher Scientific, final concentration 0.2%) and counted in a hemocytomoeter. Samples were diluted with cold NRB buffer. 90 μL aliquots containing 10 million nuclei were prepared, flash frozen in liquid nitrogen and stored at −80 °C.

For biological replicates, nuclei were isolated independently (i. e. cells were seeded and differentiated on different days), but all steps described below were performed in parallel.

Drosophila S2 nuclei were prepared and used as Spike-Ins in the PRO-Cap- and PRO-Seq experiments. Therefore, 100,000,000 Drosophila S2 cells were kindly provided by the lab of Alexander Stark. They were distributed to two tubes and centrifuged at 1000 g and 4 °C for 5 min. Cells in each tube were resuspended in 15 mL of cold 1x PBS and centrifuged at 1000 g and 4 °C for 5 min. Cells in each tube were resuspended in 1.5 mL of cold IA buffer and then pooled, incubated on ice for 3 min and centrifuged at 700 g and 4 °C for 5 min. After removing the supernatant, they were again resuspended in 2 mL of cold IA buffer, incubated on ice for 3 min and centrifuged at 700 g and 4 °C for 5 min. The supernatant was removed, nuclei were resuspended in 200 μL of cold NRB buffer and transferred to a fresh, RNase-free 1.5 mL reaction tube. Nuclei were stained with Trypan Blue Solution and counted in a hemocytometer. As nuclei tended to be lysed quickly by the Trypan Blue, they were counted immediately after adding the Trypan Blue to ensure accurate estimation of nuclei numbers. Samples were diluted with cold NRB buffer. Aliquots containing 50,000 nuclei/μL were prepared, flash frozen in liquid nitrogen and stored at −80 °C.

### Nuclear Run-On

A 2x NRO-mix was prepared containing 10 mM TRIS-HCl pH 8.0, 5 mM Mg Cl2, 1 mM DTT, 300 mM KCl, 0.05 mM Biotin-11-CTP (Biotium), 0.05 mM Biotin-11-UTP (Biotium), 0.05 mM ATP (Sigma Aldrich), 0.05 mM GTP (Sigma-Aldrich), 0.4 U/μL SUPERaseIn RNase Inhibitor (Fisher Scientific) and 1% sarkosyl. By using only two biotinylated nucleotides instead of four, we cannot achieve the single base-pair resolution of the original PRO-Seq method (Mahat *et al*., 2016), as incorporation of ATP or GTP will not lead to abortion of the run-on reaction. However, for our purposes this reduced resolution is still sufficient and with this modified protocol we can avoid including costly Biotin-ATP and Biotin-GTP in the run-on reaction. The NRO-mix was pre-warmed to 30 °C.

ESC and S2 nuclei were thawed on ice. 10 μL of S2 aliquots containing 50,0000 nuclei were added resulting in a total volume of 100 μL of ESC/S2 nuclei in NRB. To ensure identical run-on duration between different samples, for the following steps only one sample was handled at a time. 100 μL of nuclei were added to 100 μL of pre-warmed 2x NRO-mix. Samples were mixed gently by pipetting up and down 15 times and nuclear run-on was performed by incubation at 30 °C for exactly 3 min. After 90 seconds (s), samples were briefly mixed by gentle tapping. The run-on was stopped by adding 500 μL of TRI Reagent® LS (Sigma-Aldrich), samples were incubated for 5 min at room temperature and flash frozen in liquid nitrogen.

### PRO-Seq

For PRO-Seq we largely followed a previously published protocol (Mahat *et al*., 2016) with some adjustment as described below. Run-on reactions were thawed and RNA was extracted as described previously. However, during all RNA extraction steps samples were centrifuged at 20000 g and 4 °C. In addition, RNA pellets were washed with 80% ethanol and only air-dried for 2 min after carefully removing as much supernatant as possible. Moreover, when pre-washing the Streptavidin beads, all incubation steps were performed for 2 min.

Base hydrolysis was optimized and performed with 5 μL of 1 M NaOH for 20 min. In addition, SUPERaseIn RNase Inhibitor was used whenever the previously published protocol suggested to use RNase inhibitor. We also used TRI Reagent® instead of Trizol, and we used RNase-free, but not DEPC-treated water (Sigma-Aldrich).

3’-adapter ligation was performed at 16 °C overnight. For 5’ cap repair, 2.5 U of Cap-Clip^TM^ Acid Pyrophosphatase (Biozym) and its reaction buffer were used instead of TAP or RppH enzymes. After 5’ hydroxyl repair, a single RNA extraction was performed with 500 μL of TRI Reagent® and 100 μL of chloroform. 5’ adapter ligation was also performed at 16 °C overnight. For reverse transcription, the RP1 primer was used.

For PCR amplification of the libraries, we used the PCR amplification mix from the sparQ DNA Library Prep Kit. After reverse transcription, 1 μL of 35 μM forward (RPI1-10) and reverse primers (RP1) as well as 3 μL of water and 25 μL of PCR amplification mix were added to the 20 μL sample. As barcodes were introduced with the forward PCR primer, a different forward primer was used for each library to allow for multiplexing samples and including them in the same sequencing run. The number of cycles for optimal PCR amplification was estimated to be 9-14 in total as described above for the ChIP-Seq libraries.

After PCR amplification, samples were stained with SYBR® Green I nucleic acid gel stain and run on a 2.5% low melt agarose gel prepared with 0.5x TBE and run in 1x TBE for 25 min at 100 V. The part of the gel corresponding to 100-300 bp was cut and libraries were gel-extracted with the NucleoSpin^TM^ Gel and PCR Clean-up kit (Macherey-Nagel^TM^). Libraries were quantified and pooled as described above for ChIP-Seq. The size distribution of the pooled libraries was analyzed with the Agilent High Sensitivity DNA kit. To remove residual primers and adapters, an additional purification step with 1.4x AMPure XP beads was performed. After removing the supernatant containing primers and adapters, libraries were eluted from the beads and sequenced at the VBCF NGS Unit. Due to the adapter design, sequencing reads correspond to the reverse complement of the nascent RNA.

### PRO-Cap-Seq

For PRO-Cap-Seq, we largely followed the same published protocol as for PRO-Seq with the modifications described above. In addition to this, we included a buffer exchange with a P-30 column - as described in the PRO-Seq protocol - before the very first biotin-enrichment with Streptavidin beads.

We also performed 3’ adapter ligation with 2 μL of T4 RNA ligase 2, truncated K227Q (NEB) and ATP-free T4 RNA ligase buffer in a total volume of 21 μL at 16 °C overnight, as we used a 3’ DNA rather than RNA adapter.

Moreover, we chose a modified strategy for 5’ end modification. Rather than degrading 5’ mono-phosphate-containing RNAs and removing 5’ tri- and monophosphates, we decided to dephosphorylate all 5’ ends except of those protected by a 5’-cap. In an ensuing step, the 5’-cap was removed leaving behind a 5’ phosphate. This strategy ensures that 5’-adapter ligation – which requires a 5’ phosphate – only occurs on RNA molecules that had previously been capped.

Therefore, we performed biotin RNA enrichment as described before and resuspended the RNA pellet in 10 μL of RNase-free water. After denaturation for 20 s at 65 °C, RNA was stored on ice and 1 U of Shrimp Alkaline Phosphatase (NEB), 1 μL of SUPERaseIn RNase Inhibitor and 2 μL of 10xCutSmart® Buffer (NEB) were added. RNase-free water was added to a final volume of 20 μL. After incubation at 37 °C for 1 h, RNase-free water was added to a final volume of 100 μL and RNA was extracted with 500 μL TRI Reagent® and 100 μL chloroform as described previously.

The RNA pellet was resuspended in 5 μL of RNase-free water and treated with Cap-Clip^TM^ enzyme as described above for the PRO-Seq. RNA was extracted with 500 μL TRI Reagent® and 100 μL of chloroform once more. 1 μL of 5’ RNA adapter (50 μM) was diluted in 4 μL of RNase-free water and the RNA pellet was dissolved in this RNA-adapter dilution. After denaturation at 65 °C for 20 s, 2.2 μL of 10x T4 RNA ligase buffer (NEB), 6 μL 50%PEG 8000, 10 mM ATP, 1 μL SUPERaseIn RNase Inhibitor, 1 μL T4 RNA ligase 1 (NEB, 10 U) and RNase-free water (to a total volume of 22 μL) were added. 5’ adapter ligation was performed at 16 °C overnight.

Biotin-RNA enrichment and reverse transcription were performed as described previously. However, for reverse transcription different primers were used for every sample (RPIC1-4), as barcodes for multiplexing were already introduced in this step.

PCR amplification was performed with the KAPA HiFi Real-Time PCR library amplification kit (Roche). Therefore, 1 μL of 35 μM forward (RPC1) and reverse primer (RPIC1-4) as well as 3 μL of water and 25 μL of 2x KAPA HiFi amplification mix were added to the 20 μL of cDNA. PCR amplification was performed according to the standard protocol in a qPCR machine. This allowed to measure both fluorescence of the standards included in the KAPA kit and fluorescence of the amplified libraries, and thus to monitor the amplification status. For each library, amplification was stopped shortly after the curve depicting the relative fluorescence units for each cycle started to show exponential growth.

PRO-Cap-Seq libraries were run on a 2.5% low-melt agarose gel and gel-extracted as described above. Libraries were quantified, pooled and the size distribution of the pooled libraries was analyzed as described above. Sequencing was performed at the VBCF NGS Unit.

### Data analysis

PRO-Cap-Seq libraries were sequenced to a depth of 22-30 million reads while PRO-Seq libraries were sequenced to a depth of 30-60 million reads (both: single-end, 50 bp). For both PRO-Seq and PRO-Cap-Seq we used adapters containing random nucleotides of 4 (PRO-Seq 5’ and PRO-Cap-Seq 3’ adapter), 8 (PRO-Seq, 3’ adapter) or 10 bp (PRO-Cap-Seq, 5’ adapter) length. This allowed us to distinguish between identical reads that are PCR duplicates – those should have the exact same random nucleotides as they are amplified from the same molecule – and identical reads that originate from different RNA molecules with the same sequence – for those it is highly unlikely to have the exact same random nucleotides in the adapters.

As PRO-Seq libraries were sequenced from the 3’ end, and PRO-Cap-Seq libraries were sequenced from the 5’ end, the first eight/ten nucleotides of every unprocessed read correspond to the random nucleotides. Therefore, we removed PCR duplicates by simply removing all unprocessed reads with exact identical sequence. For this purpose, we used the Clumpify tool from BBTools version 37.20 (https://github.com/BioInfoTools/BBMap/blob/master/sh/clumpify.sh). Specified adapters were removed with Trim Galore Version 0.5.0 (https://github.com/FelixKrueger/TrimGalore); in addition to this, the first eight (PRO-Seq)/ten (PRO-Cap-Seq) nucleotides of every read were trimmed as those correspond to the random nucleotides and would interfere with genome alignment later on. We also trimmed the last four nucleotides of every read, as those might potentially represent the random nucleotides introduced by the 5’ (PRO-Seq)/3’ (PRO-Cap-Seq) adapter.

As the reads in both PRO- and PRO-Cap-Seq libraries were a mixture of nascent transcripts from ESCs and S2 Spike-Ins, we generated a genome assembly merged from the mm10 assembly of the mouse genome and a current release of the Drosophila melanogaster genome downloaded from Flybase (Thurmond *et al*., 2019) for alignment. We preferred this strategy over first aligning to the mouse and then to the Drosophila genome, as with our strategy we could exclude reads that mapped to both genomes and for which we could not be sure, whether they originate from our actual samples or from the Spike-Ins. With the alternative strategy, all of those reads would have been assigned to the ESC samples. For KO and KI cell lines, custom mm10 genomes carrying the corresponding genetic modifications were assembled with the help of the *reform* tool (https://github.com/gencorefacility/reform) and then merged with the Drosophila genome.

We performed alignment with Bowtie 2 Version 2.3.4.3 (Langmead & Salzberg, 2012). SAMtools 1.5 (H. Li *et al*., 2009) was used to convert the resulting sam files to bam files, to sort and index the bam files as well as for extracting uniquely mapping reads. We also used SAMtools 1.5 to separate bam files with uniquely mapping reads into two files with reads mapping to mouse and Drosophila genome respectively, and to split the resulting files by which strand reads were mapping to. In case of the PRO-Seq libraries, we accounted for the fact that sequencing reads correspond to the reverse complement of the nascent RNA i. e. reads mapping to the minus strand originated from transcripts with the sequence of the plus strand and *vice versa*.

For PRO-Cap-Seq libraries, we also used the GATK ClipReads version 4.0.1.2 (McKenna *et al*., 2010) function to trim aligned reads to the very first nucleotide; this is the nucleotide at which transcription had been initiated. We decided not to do the same for the PRO-Seq libraries, because, as mentioned above, we used only two biotinylated nucleotides for the Run-On and thus did not have the single-bp resolution required for an unbiased analysis of which nucleotide had been incorporated last during transcription.

For visualization, bam files containing uniquely mapping reads were converted into bedgraph files with bedtools version 2.28.0 (Quinlan & Hall, 2010) while normalizing for sequencing depth of the respective Spike-In. Bedgraph files were then converted to bigWig files using the bedGraphToBigWig ((https://github.com/sccallahan/bedGraph2bigWig)) tool. BigWig files were visualized with the UCSC genome browser (Kent *et al*., 2002).

For quantitative analysis, we generated gtf files containing the genomic features of interest (such as the different enhancers at the locus), and counted reads mapping to these features with the featureCounts function of the Rsubread package (version 1.5.3) (Liao *et al*., 2019). For enhancers, we counted reads within a 1500 bp window centered on the p300 peak. Only for the PE element, we used a smaller 800 bp window to minimize the effect of reads originating from the nearby promoter. The 800 bp correspond to the size of the element that had been reintroduced for generating the PE KI cell lines.

To calculate the travel ratio, we counted reads in the gene body (all reads mapping between start of exon two and end of exon three), and divided them by the reads counted in a 350 bp window around the TSS (as defined by PRO-Cap-Seq signal). We manually normalized to sequencing depth of the Spike-Ins and generated graphs with the ggplot2-3.3.0 package (Wickham, 2016)

## DNA oligonucleotide sequences

**Table 1:**
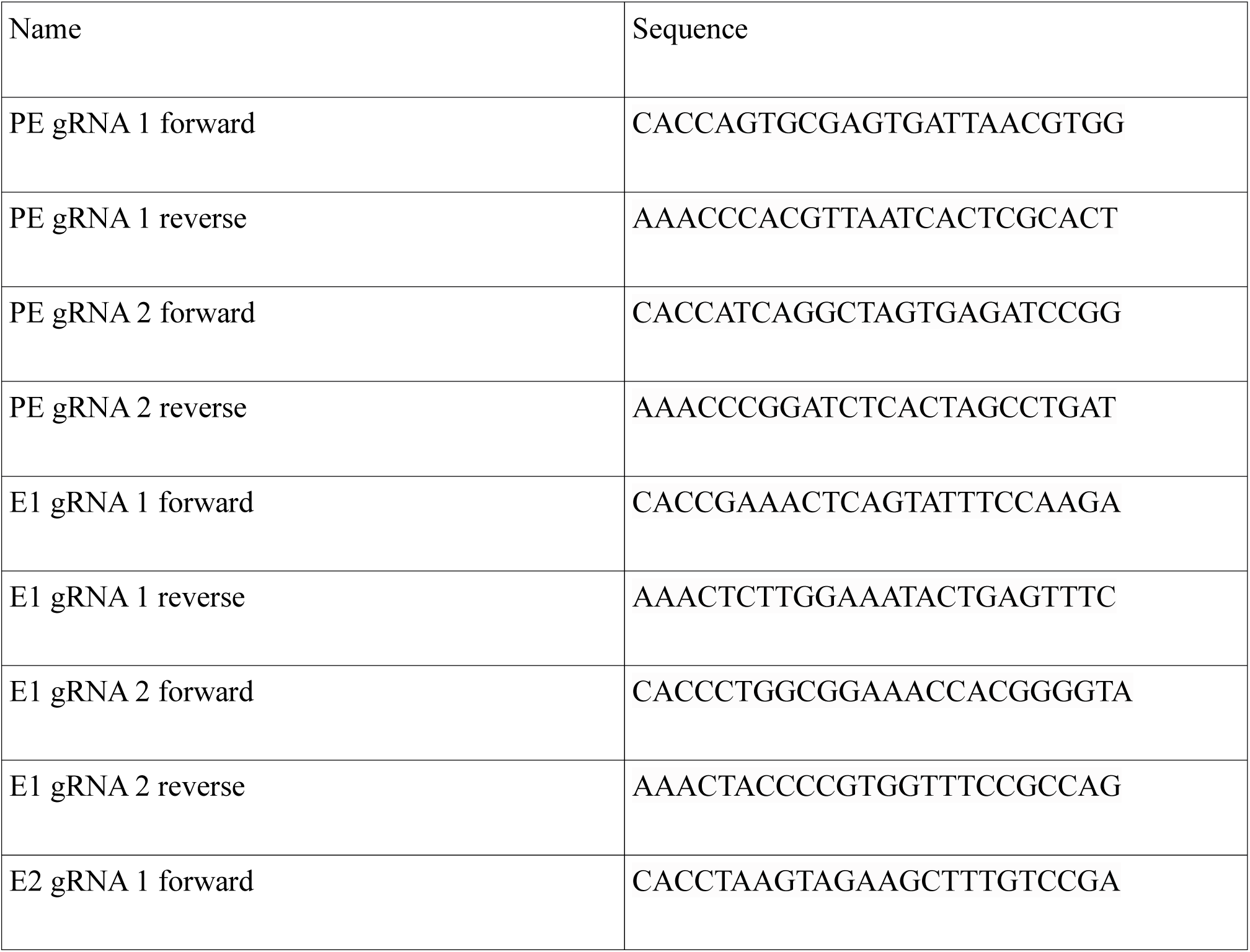
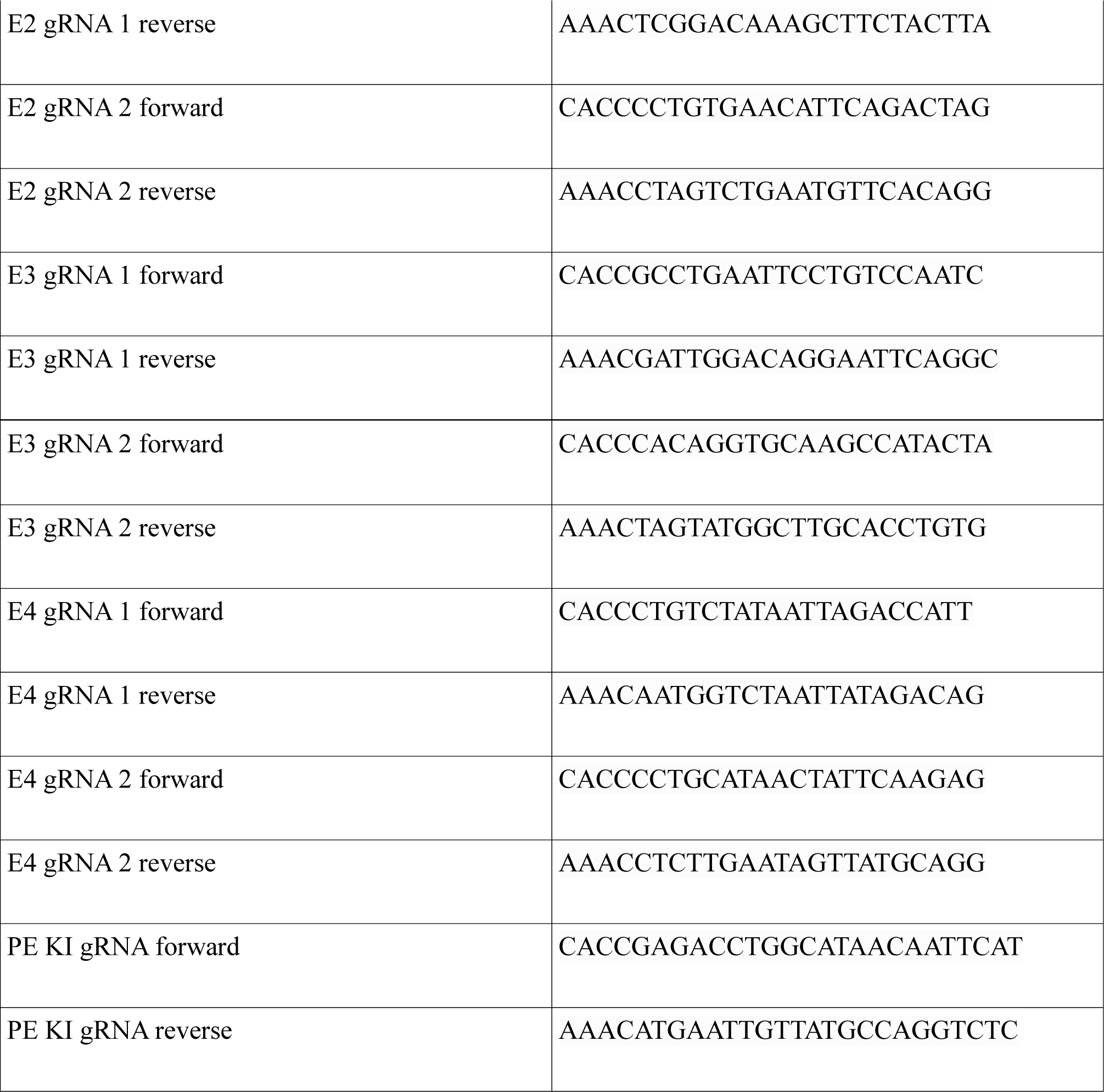
gRNAs

**Table 2:**
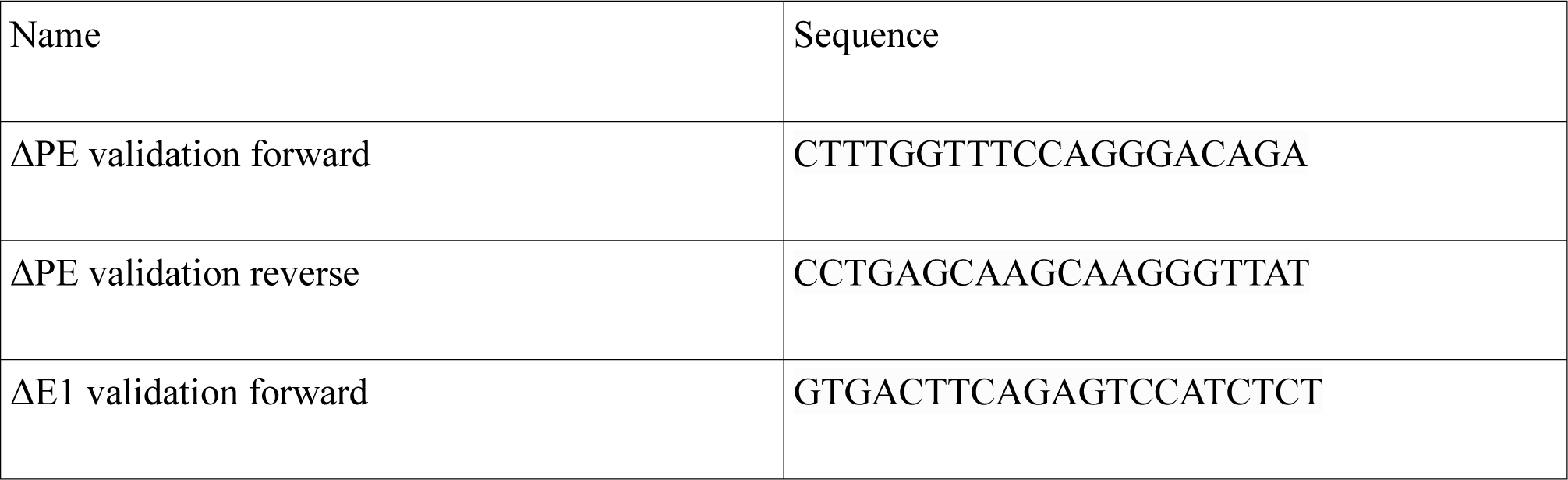
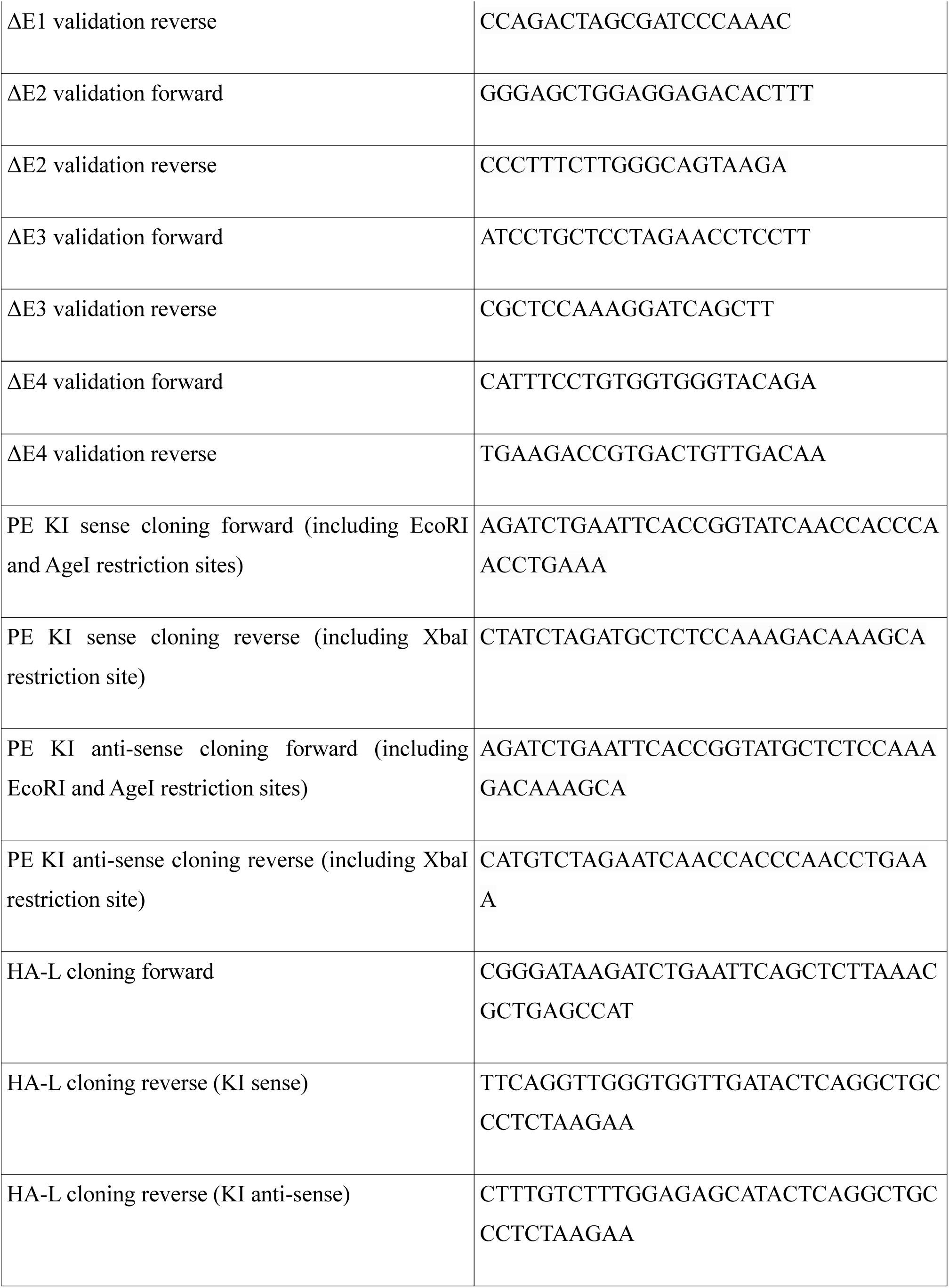
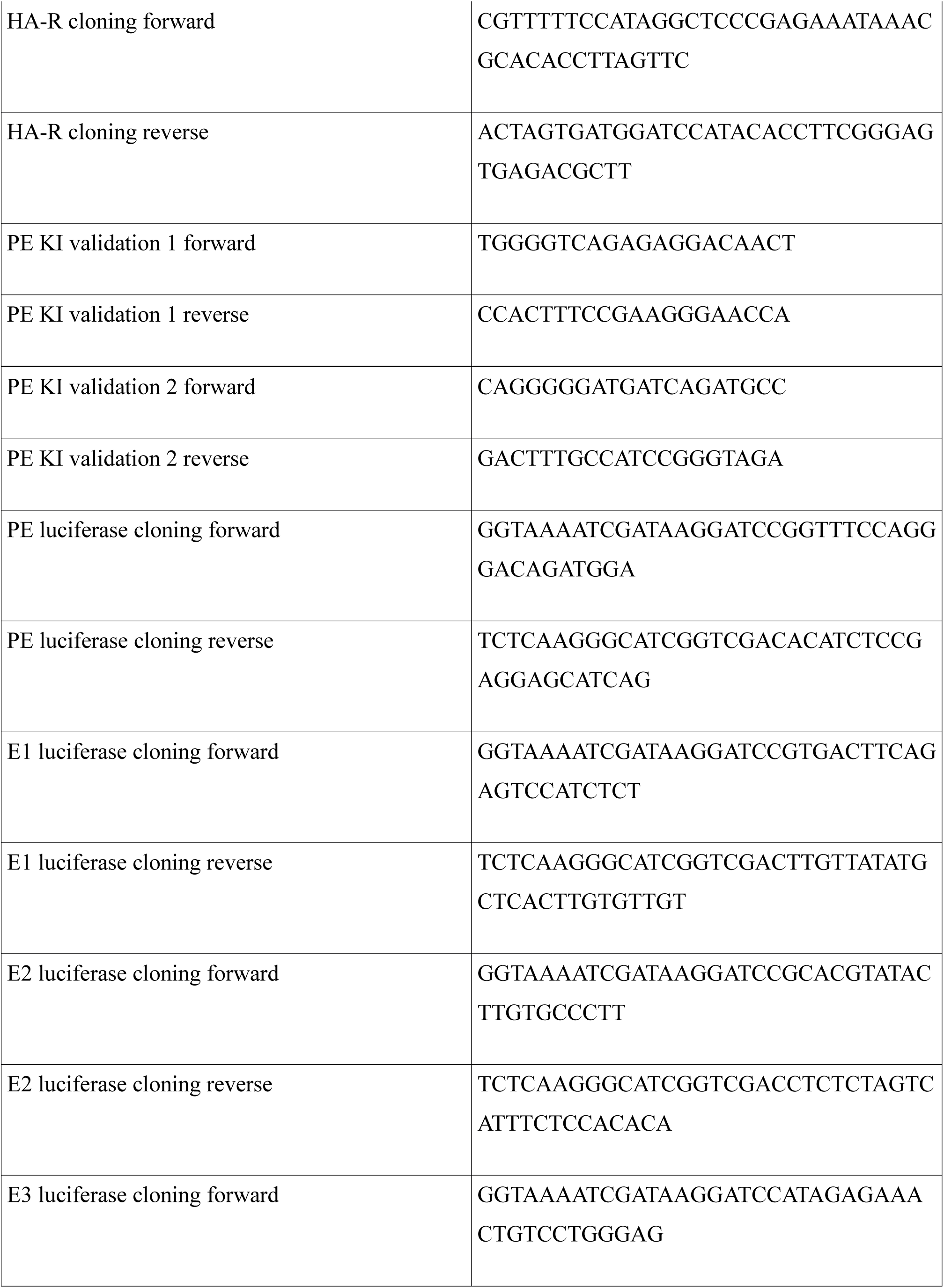
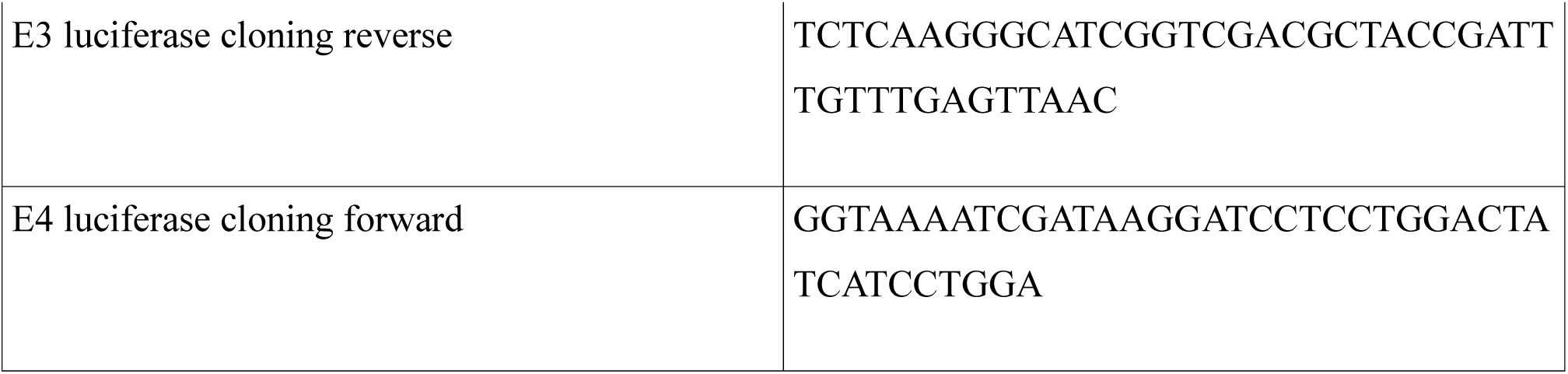
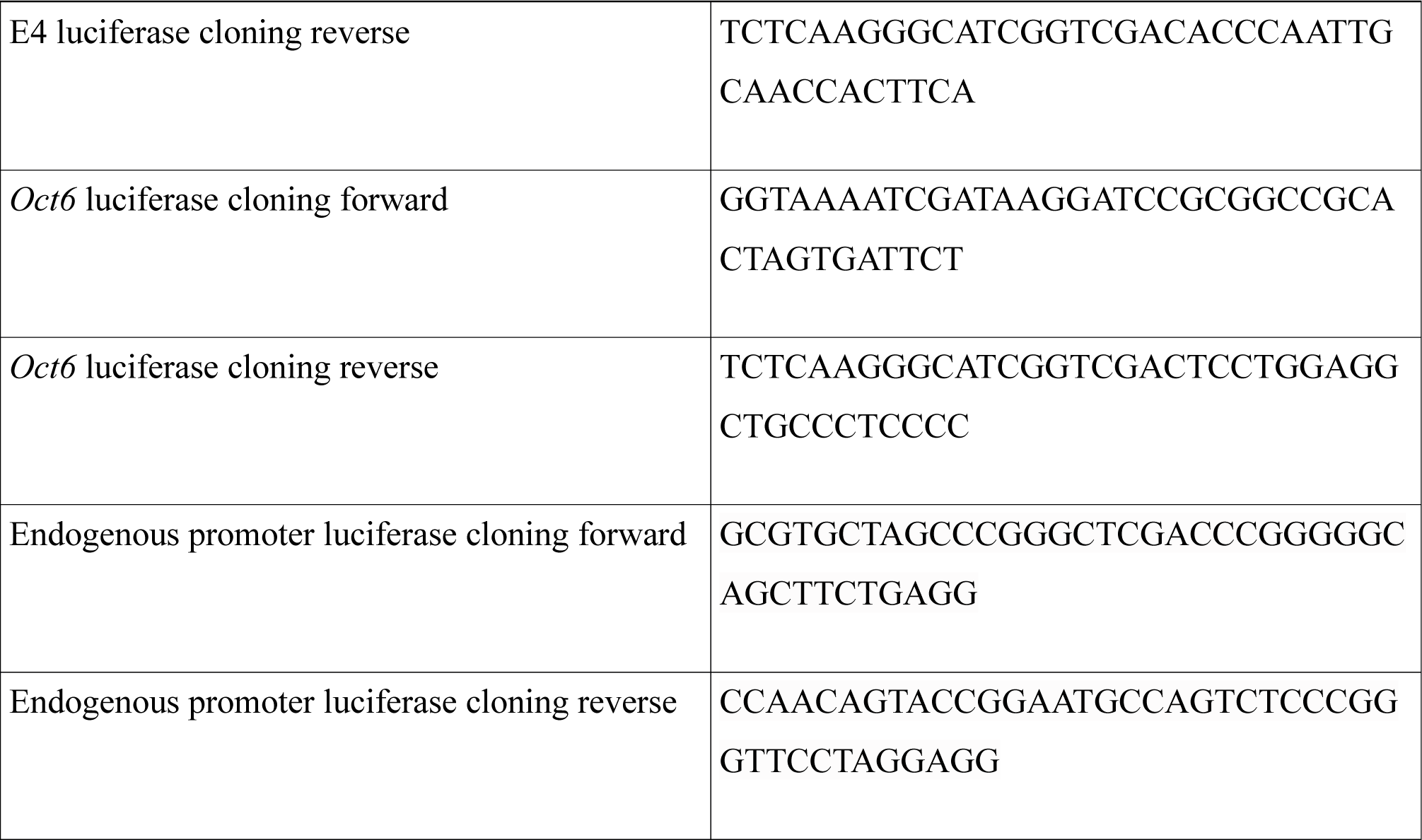
PCR primers

**Table 3:**
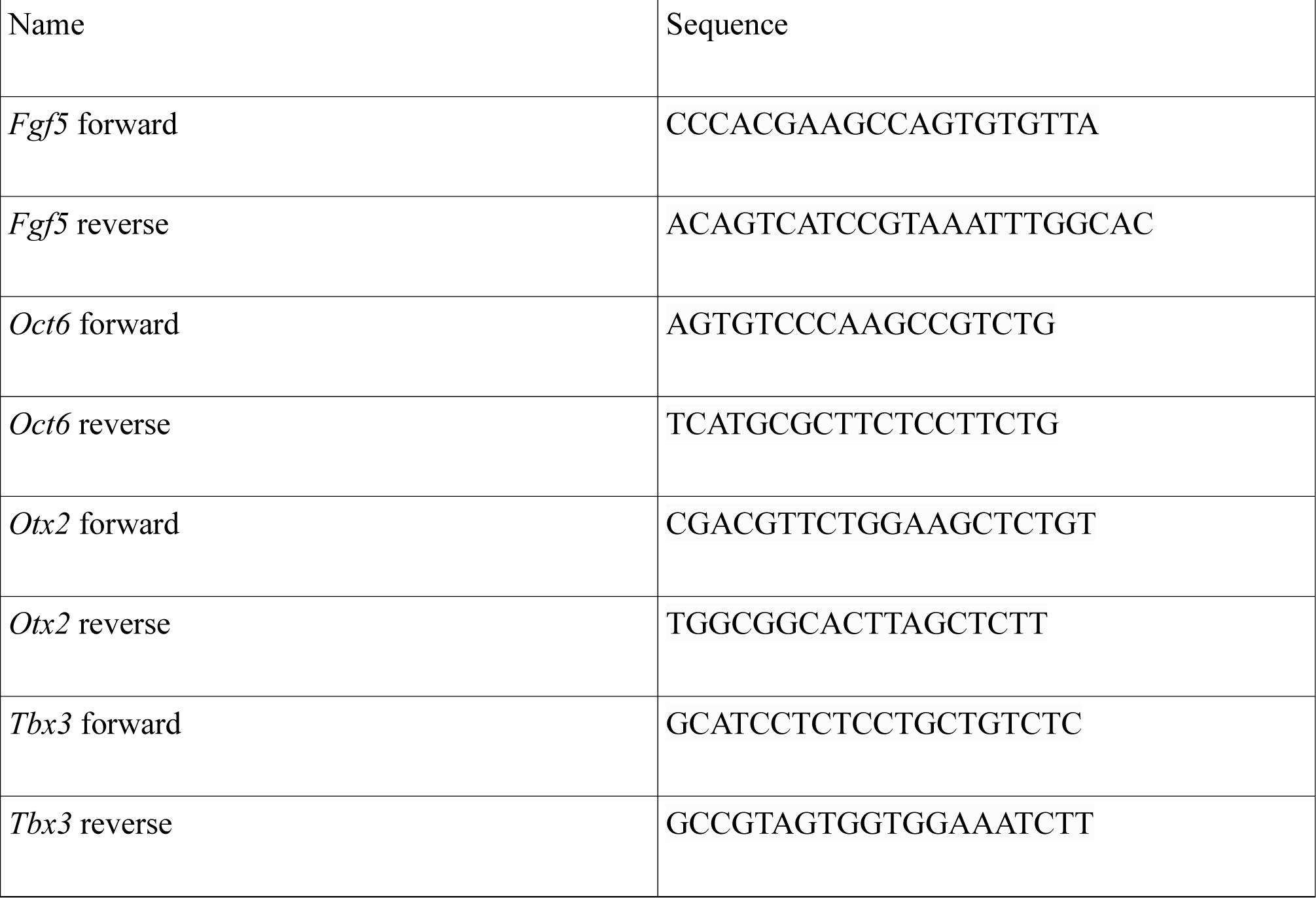
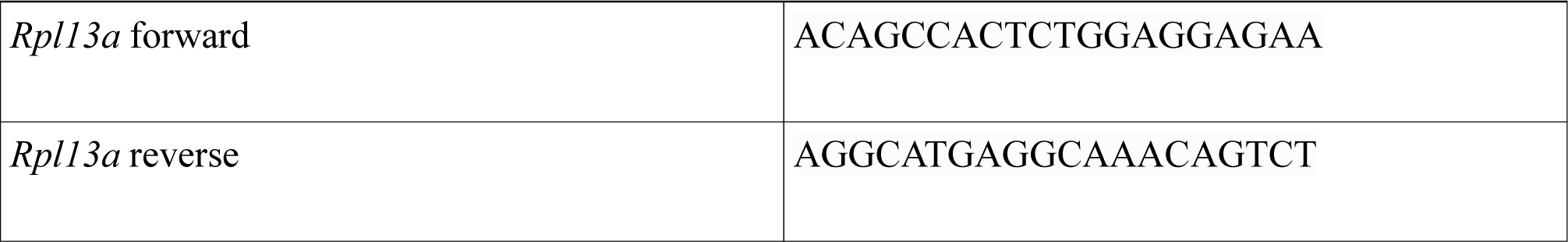
RT-qPCR primers

**Table 4:**
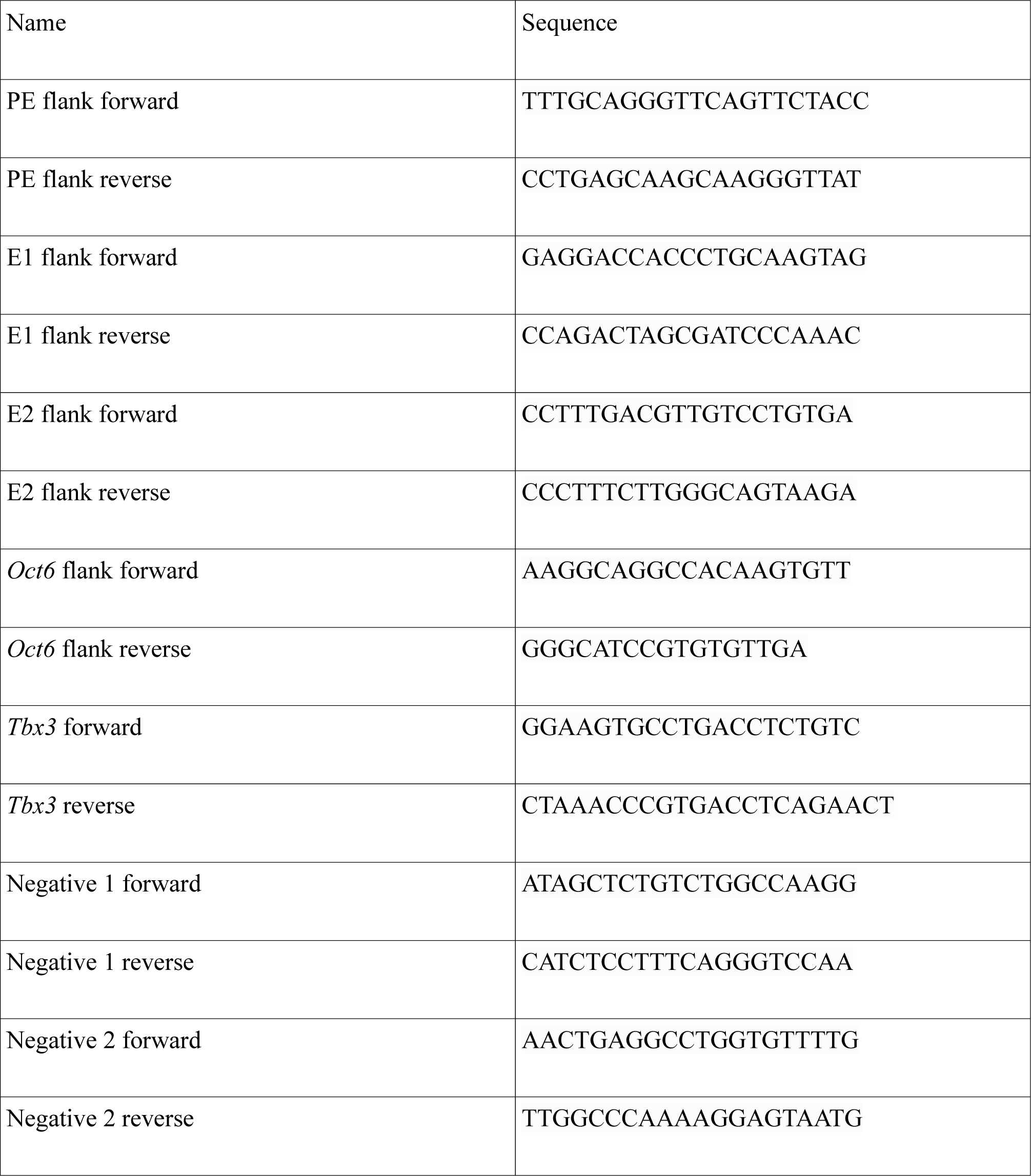
ChIP-qPCR primers

**Table 5:**
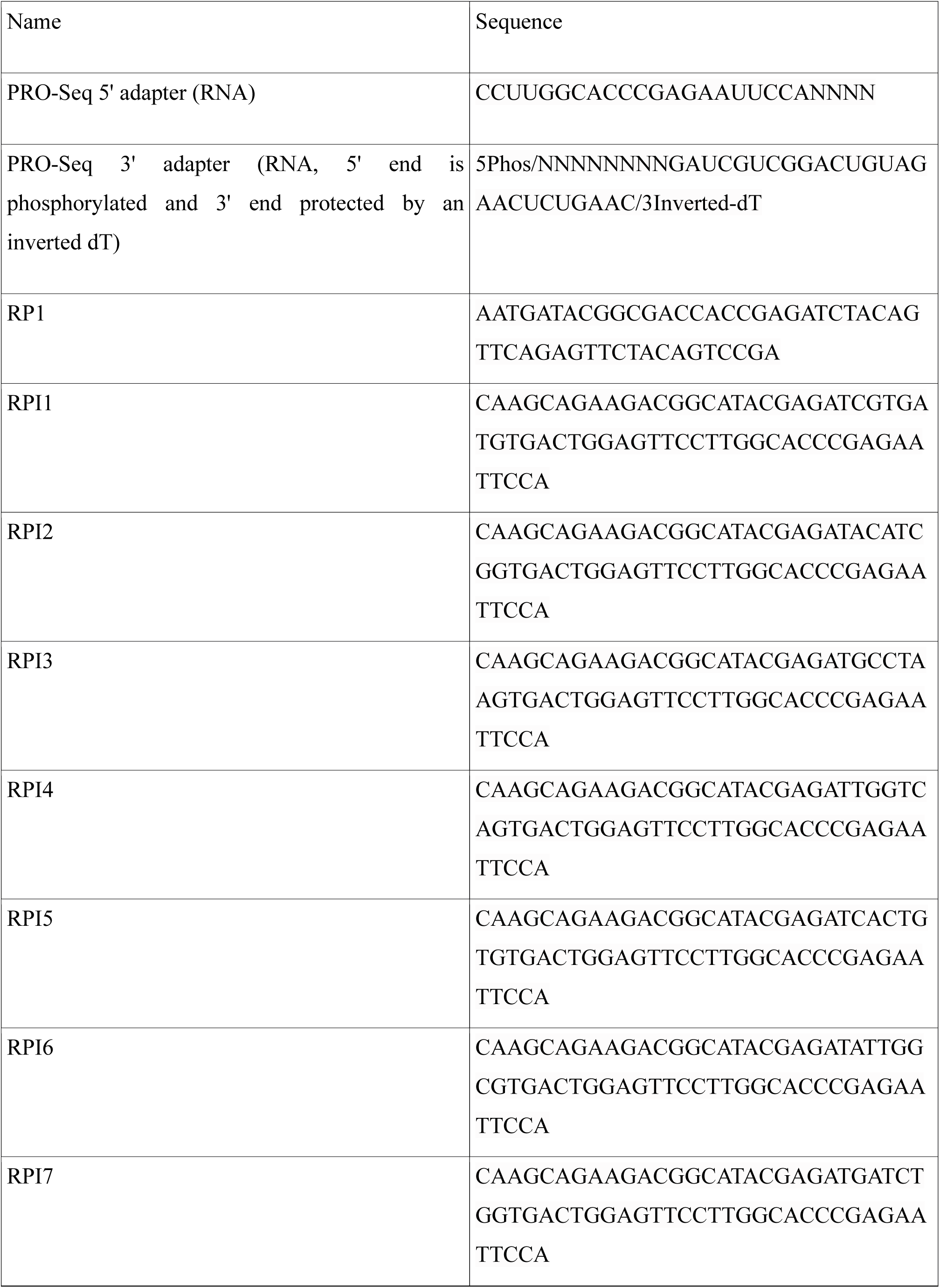
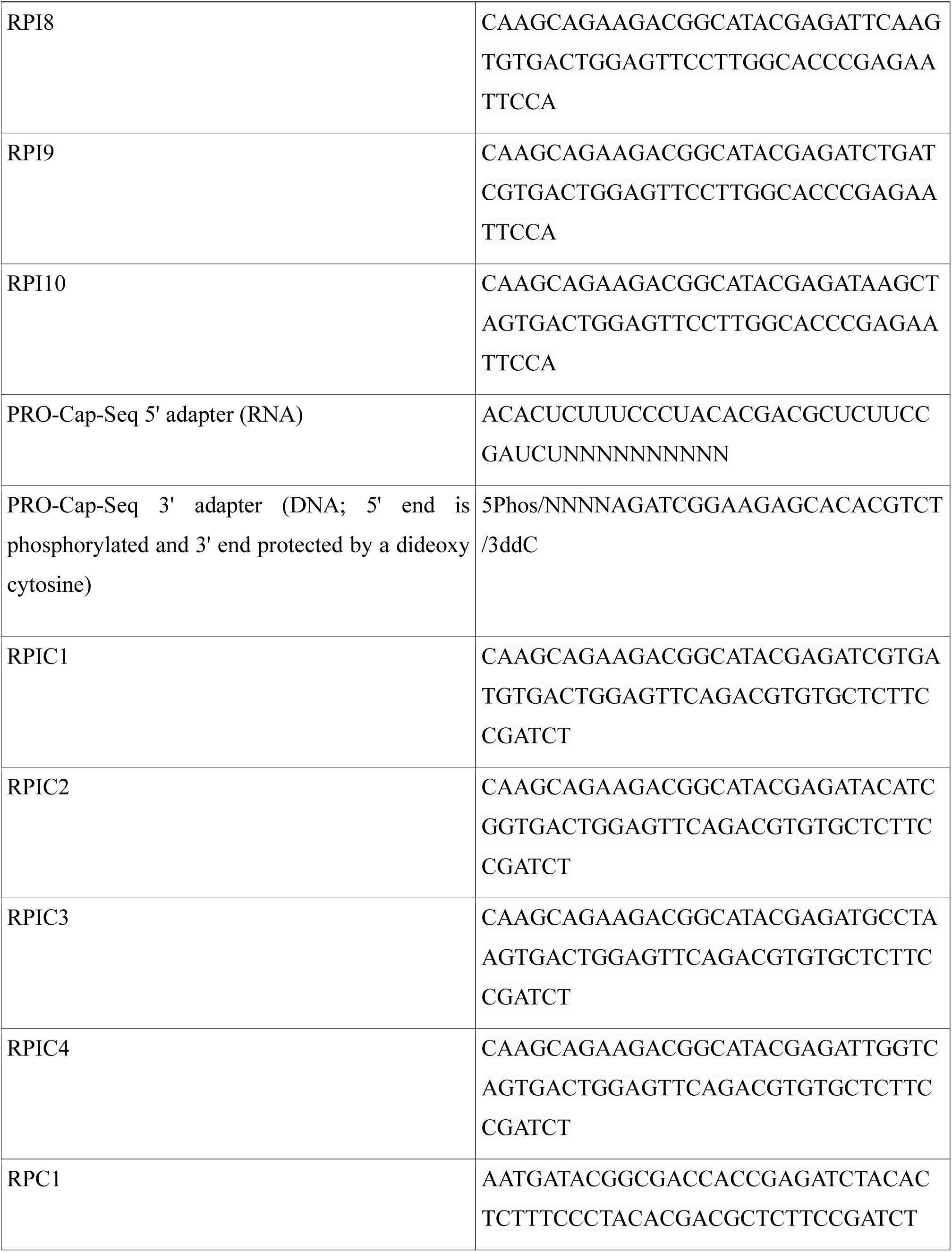
PRO- and PRO-Cap-Seq primers and adapters

---

**Supplemental Figure S1:**
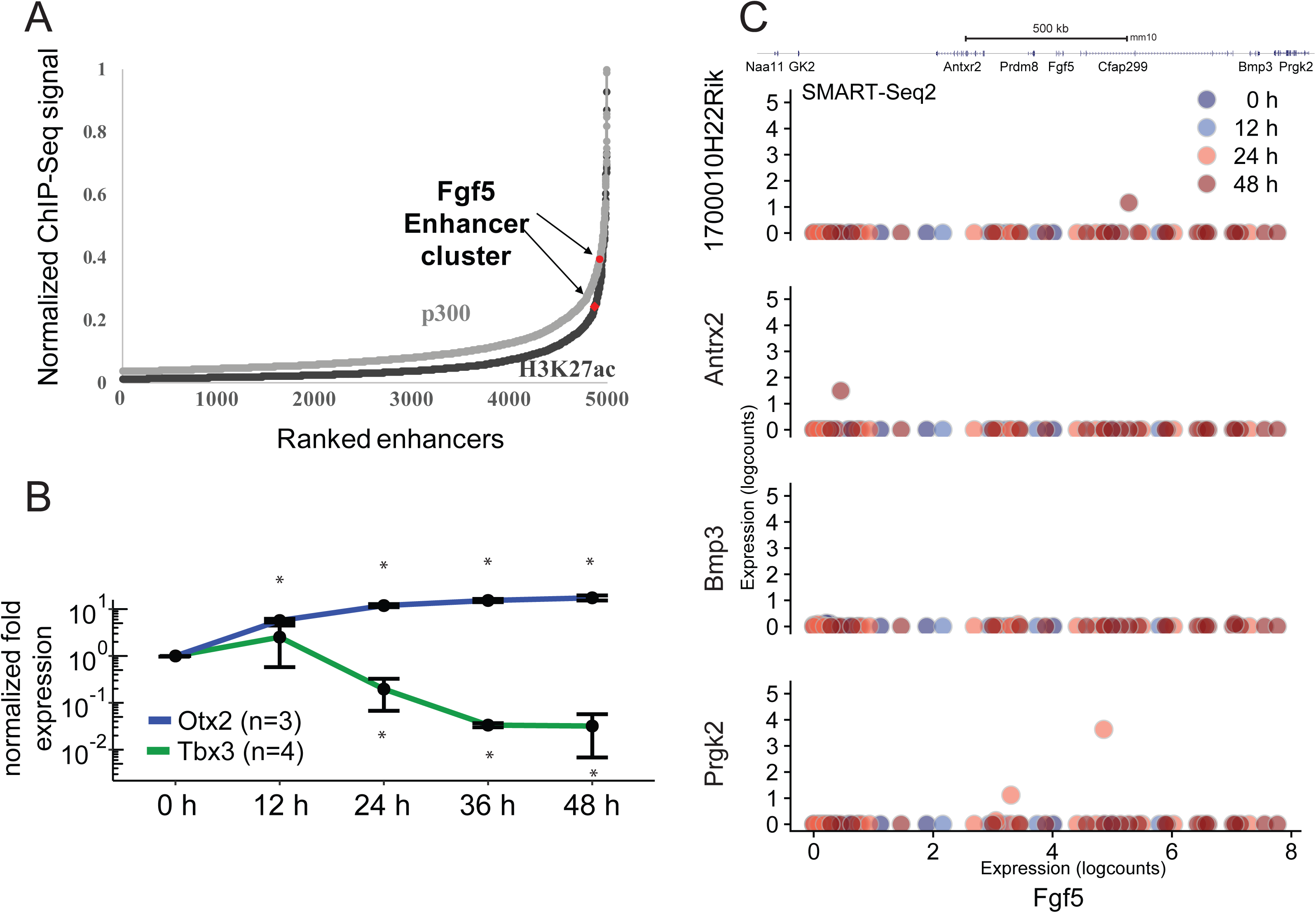
The *Fgf5* locus as a model to study collaborative enhancer action. (A) ROSE algorithm (Whyte *et al*., 2013) analysis of the EpiLC enhancer landscape. Enhancers were defined based on H3K4me1 and H3K27ac ChIP-Seq signal and enhancers within a 12.5 kb window were stitched together. The resulting enhancer clusters were ranked by p300 or H3K27ac ChIP-Seq signal. The *Fgf5* enhancer cluster is marked in red. (B) *Otx2* and *Tbx3* expression in WT cells along an ESC to EpiLC differentiation time course as determined by RT-qPCR. Expression values are normalized to *Rpl13*a and to the 0 h time point within each independent biological replicate. Mean values of n=3 (*Otx2*) or n=4 (*Tbx3*) biological replicates are shown. Error bars correspond to one standard deviation in each direction. Time points with significantly higher (*Otx2*) or lower (*Tbx3*) expression (one-sided Welch Two sample t-test) compared to 0 h are marked by stars. (C) SMART-Seq2 single-cell expression data of genes surrounding *Fgf5* at 0, 12, 24 and 48 h of ESC to EpiLC differentiation. Normalized log counts of the respective gene are plotted against normalized log counts of *Fgf5* in the same cell.

**Supplemental Figure S2:**
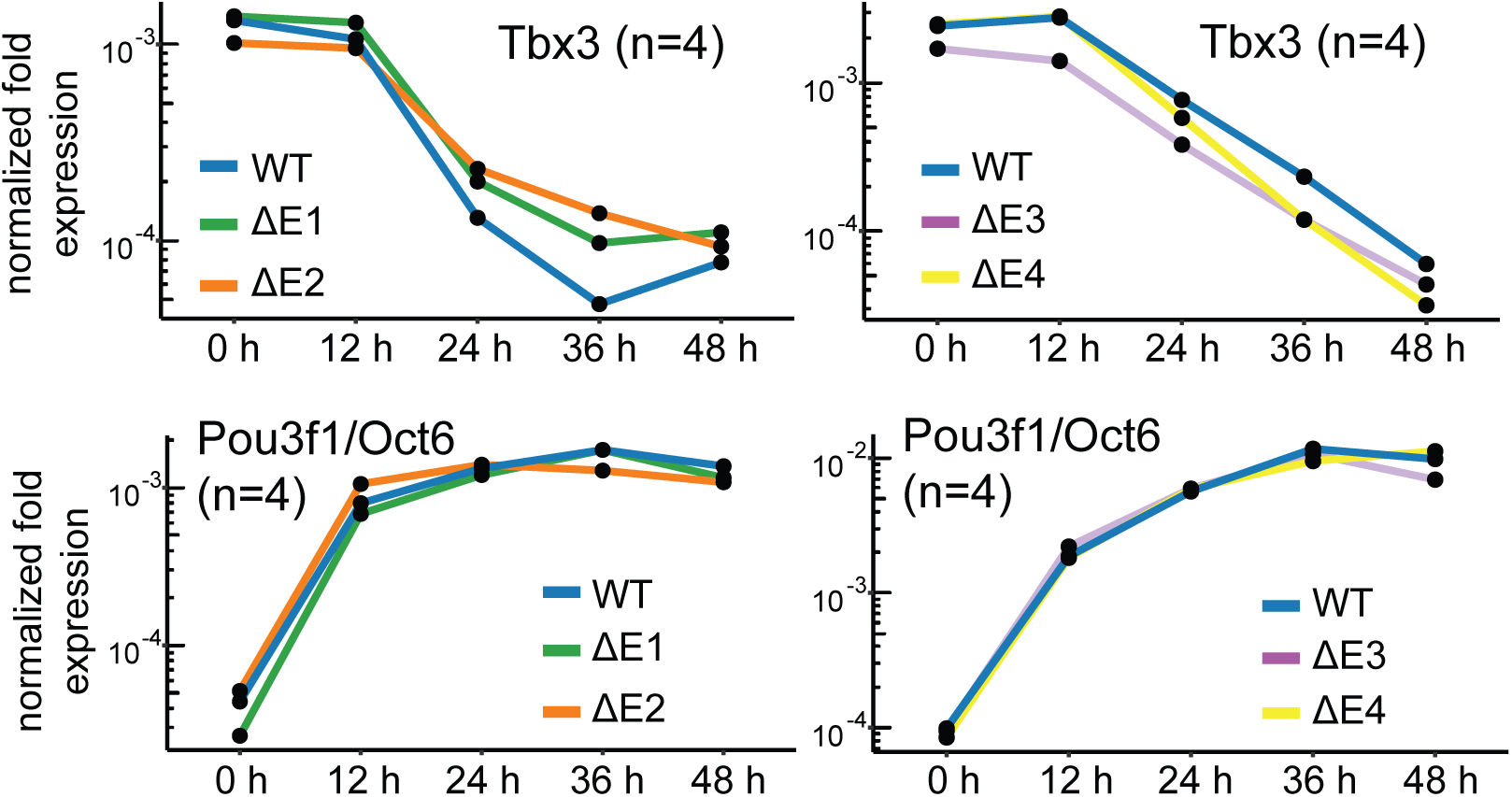
The intergenic enhancers E1-E4 mediate induction of *Fgf5* upon differentiation. (A) *Tbx3* and *Pou3f1*/*Oct6* expression in WT, ΔE1, ΔE2, ΔE3 and ΔE4 cells along an ESC to EpiLC differentiation time course as determined by RT-qPCR. Expression values are normalized to *Rpl13a*. Mean values of n=4 biological replicates are shown.

**Supplemental Figure S3:**
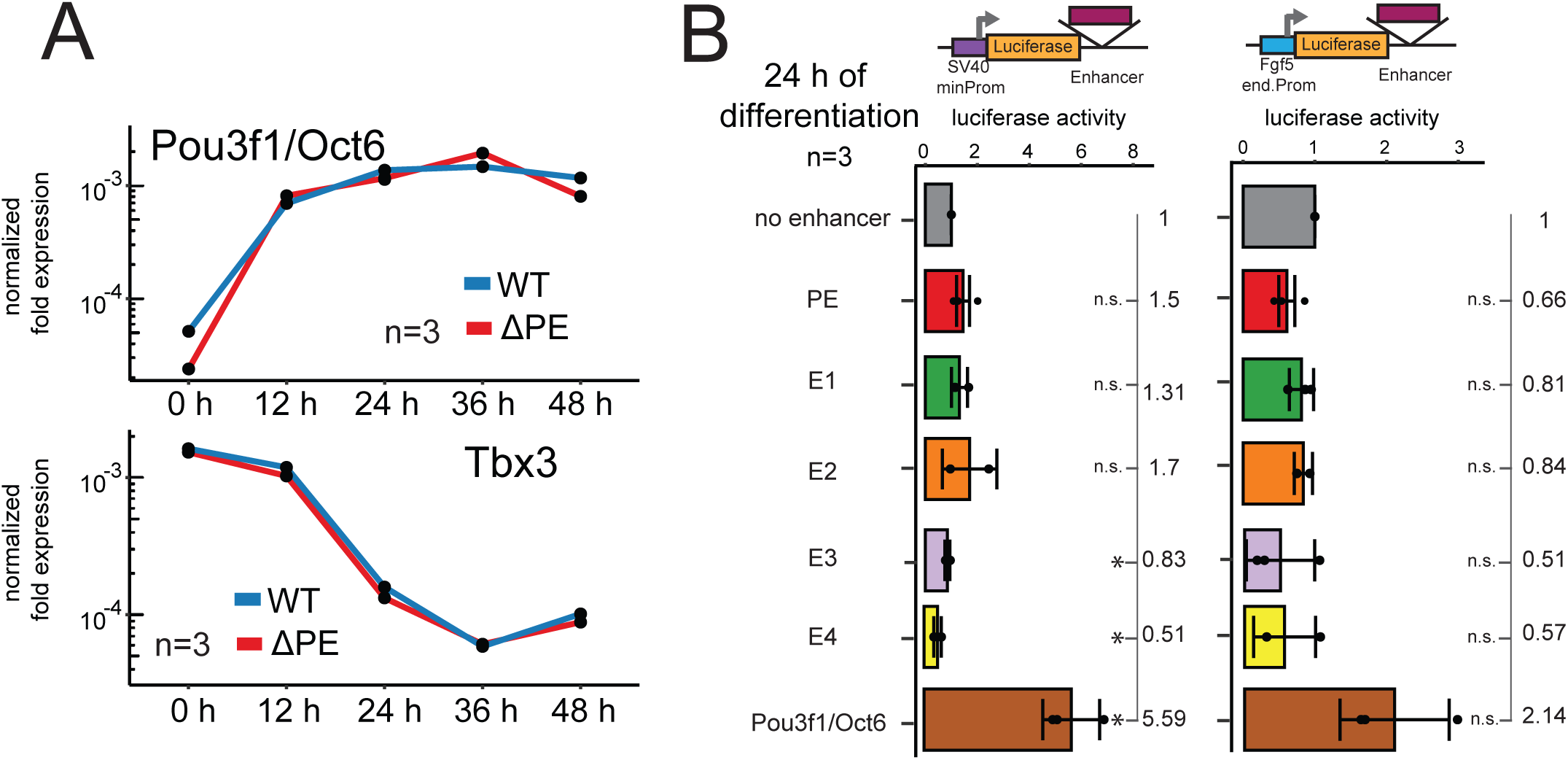
PE amplifies *Fgf5* expression levels at every time point, yet has little canonical enhancer activity in luciferase assays. (A) *Tbx3* and *Pou3f1*/*Oct6* expression in WT and ΔPE cells along an ESC to EpiLC differentiation time course as determined by RT-qPCR. Expression values are normalized to *Rpl13a*. Mean values of n=3 biological replicates are shown. (B) Luciferase assays with the respective enhancer downstream of the luciferase gene under the control of a minimal SV40 promoter (left) or under the control of the endogenous *Fgf5* promoter (right) at 24 h of differentiation. Luciferase activity is normalized first for transfection efficiency as well as plasmid size, and then to the no enhancer control within each independent biological replicate. Mean values of n=3 biological replicates are shown. Normalized values for each replicate are shown as dots. Error bars correspond to one standard deviation in each direction. Enhancers with statistically significant differences (two-sided Welch Two sample t-test) compared to the no enhancer control are marked by stars.

**Supplemental Figure S4:**
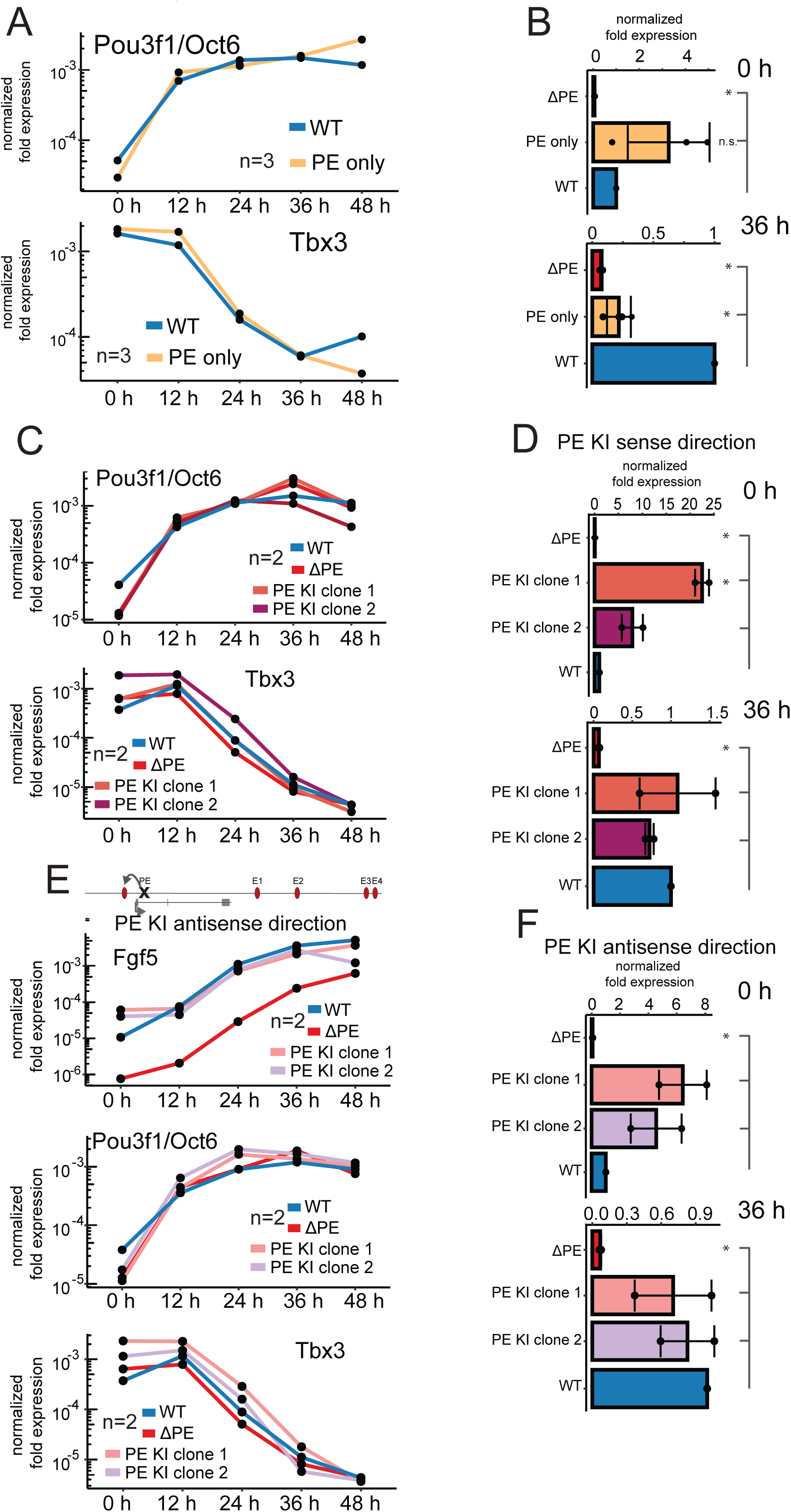
PE and E1-E4 regulate *Fgf5* transcription in super-additive fashion. (A) *Tbx3* and *Pou3f1*/*Oct6* expression in WT and PE only (individual deletion of E1 through E4) cells along an ESC to EpiLC differentiation time course as determined by RT-qPCR. Expression values are normalized to *Rpl13a*. Mean values of n=3 biological replicates are shown. (B) *Fgf5* expression in WT and PE only (individual deletion of E1 through E4) cells at 0 and 36 h of ESC to EpiLC differentiation as determined by RT-qPCR with intron-spanning primers. Expression values are normalized to *Rpl13a* and to the WT cell line at the same time point within each biological replicate. Mean values of n=3 biological replicates are shown. Normalized values for each replicate are shown as dots. Error bars correspond to one standard deviation in each direction. Cell lines with statistically lower or higher expression (one-sided Welch Two sample t-test) compared to WT at that time point are marked by stars. (C) *Tbx3* and *Pou3f1*/*Oct6* expression in WT, ΔPE and PE KI (PE integrated in sense direction) cells along an ESC to EpiLC differentiation time course as determined by RT-qPCR. Expression values are normalized to *Rpl13a*. Mean values of n=2 biological replicates are shown. (D) *Fgf5* expression in WT, ΔPE and PE KI (PE integrated in sense direction) cells at 0 and 36 h of differentiation as determined by RT-qPCR with intron-spanning primers. Expression values are normalized to *Rpl13a* and to the WT cell line at the same time point within each biological replicate. Mean values of n=2 biological replicates are shown. Normalized values for each replicate are shown as dots. Error bars correspond to one standard deviation in each direction. Cell lines with statistically lower or higher expression (one-sided Welch Two sample t-test) compared to WT at that time point are marked by stars. (E) *Fgf5*, *Tbx3* and *Pou3f1*/*Oct6* expression in WT, ΔPE and PE KI (PE integrated in anti-sense direction) cells along an ESC to EpiLC differentiation time course as determined by RT-qPCR. Expression values are normalized to *Rpl13a*. Mean values of n=2 biological replicates are shown. (F) *Fgf5* expression in WT, ΔPE and PE KI (PE integrated in anti-sense direction) cells at 0 and 36h of ESC to EpiLC differentiation as determined by RT-qPCR with intron-spanning primers. Expression values are normalized to *Rpl13a* and to the WT cell line at the same time point within each biological replicate. Mean values of n=2 biological replicates are shown. Normalized values for each replicate are shown as dots. Error bars correspond to one standard deviation in each direction. Cell lines with statistically lower expression (one-sided Welch Two sample t-test) compared to WT at that time point are marked by stars.

**Supplemental Figure S5:**
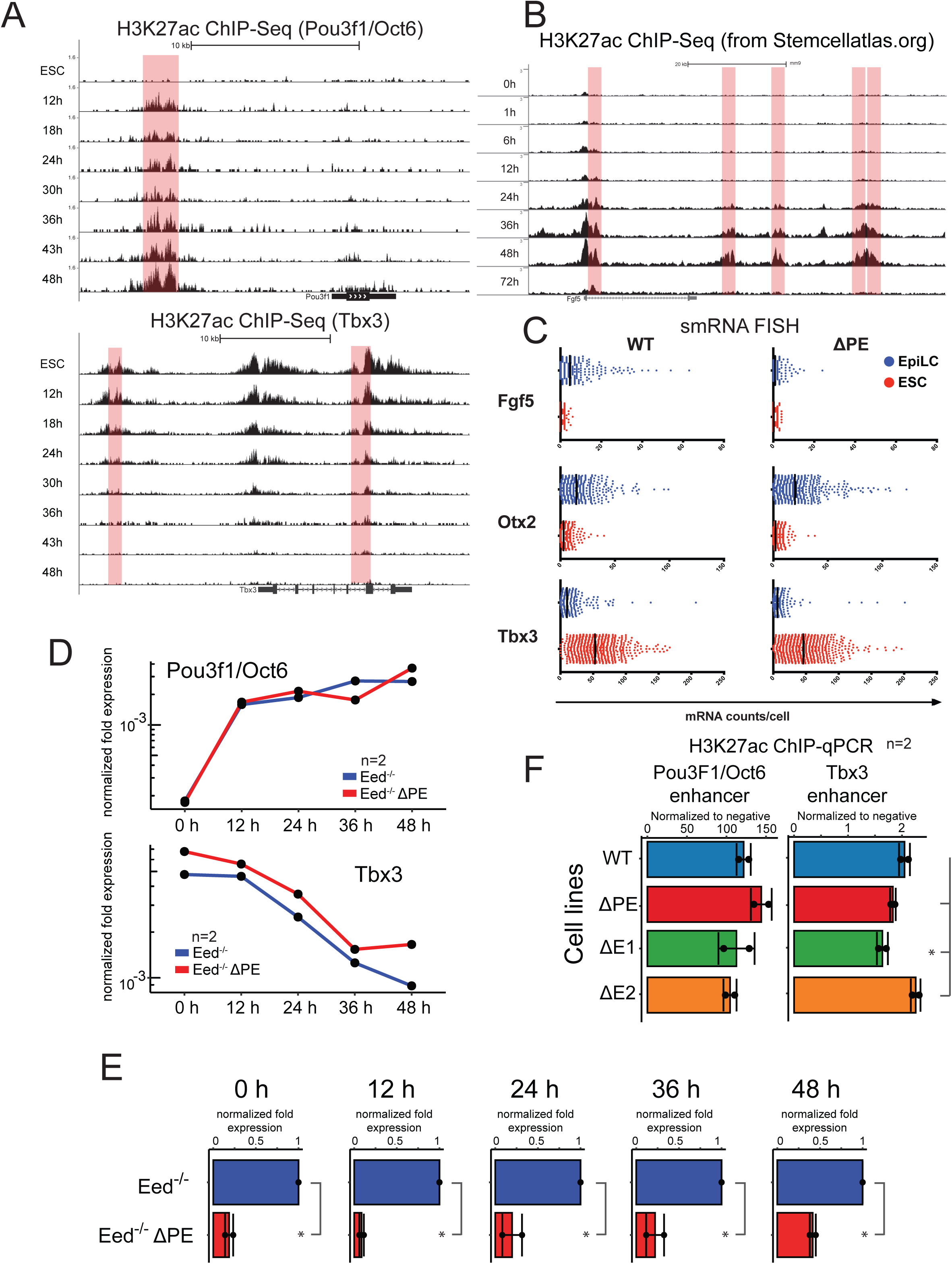
PE is not activated earlier than E1-E4 and does not primarily function by removing H3K27me3 from the *Fgf5* promoter or by facilitating activation of the intergenic enhancers E1-E4. (A) H3K27ac ChIP-Seq signal (normalized for sequencing depth) at the *Pou3f1*/*Oct6* and the *Tbx3* locus along a differentiation time course with fixed scale bar. (B) H3K27ac ChIP-Seq signal from Yang *et al*., 2019 at the *Fgf5* locus along a differentiation time course with fixed scale bar. (C) mRNA counts per cell in WT and ΔPE ESCs and EpiLCs (differentiated for 36 h) for *Fgf5*, *Otx2* and *Tbx3* as determined by ViewRNA smRNA FISH. (D) *Tbx3* and *Pou3F1*/*Oct6* expression in Eed-/- and Eed-/-ΔPE cells along an ESC to EpiLC differentiation time course as determined by RT-qPCR. Expression values are normalized to *Rpl13a*. Mean values of n=2 biological replicates are shown. (E) *Fgf5* expression in Eed-/- and Eed-/-ΔPE cells at each time point of ESC to EpiLC differentiation as determined by RT-qPCR with intron-spanning primers. Expression values are normalized to *Rpl13a* and to the Eed-/-cell line at the same time point within each biological replicate. Mean values of n=2 biological replicates are shown. Normalized values for each replicate are shown as dots. Error bars correspond to one standard deviation in each direction. Cell lines with statistically lower expression (one-sided Welch Two sample t-test) compared to Eed-/- at that time point are marked by stars. (F) H3K27ac ChIP-qPCR signal flanking the *Pou3F1*/*Oct6* and *Tbx3* enhancers in WT, ΔPE, ΔE1 and ΔE2 cells at 40 h of differentiation. Input enrichment was calculated and then normalized within each individual sample to two genomic regions known not to be marked by H3K27ac by previous ChIP-Seq studies (Buecker *et al*., 2014). Mean values of n=2 biological replicates are shown. Normalized values for each replicate are shown as dots. Error bars correspond to one standard deviation in each direction. Cell lines with statistically lower signal (one-sided Welch Two sample t-test) compared to WT at that time point are marked by stars.

**Supplemental Figure S6:**
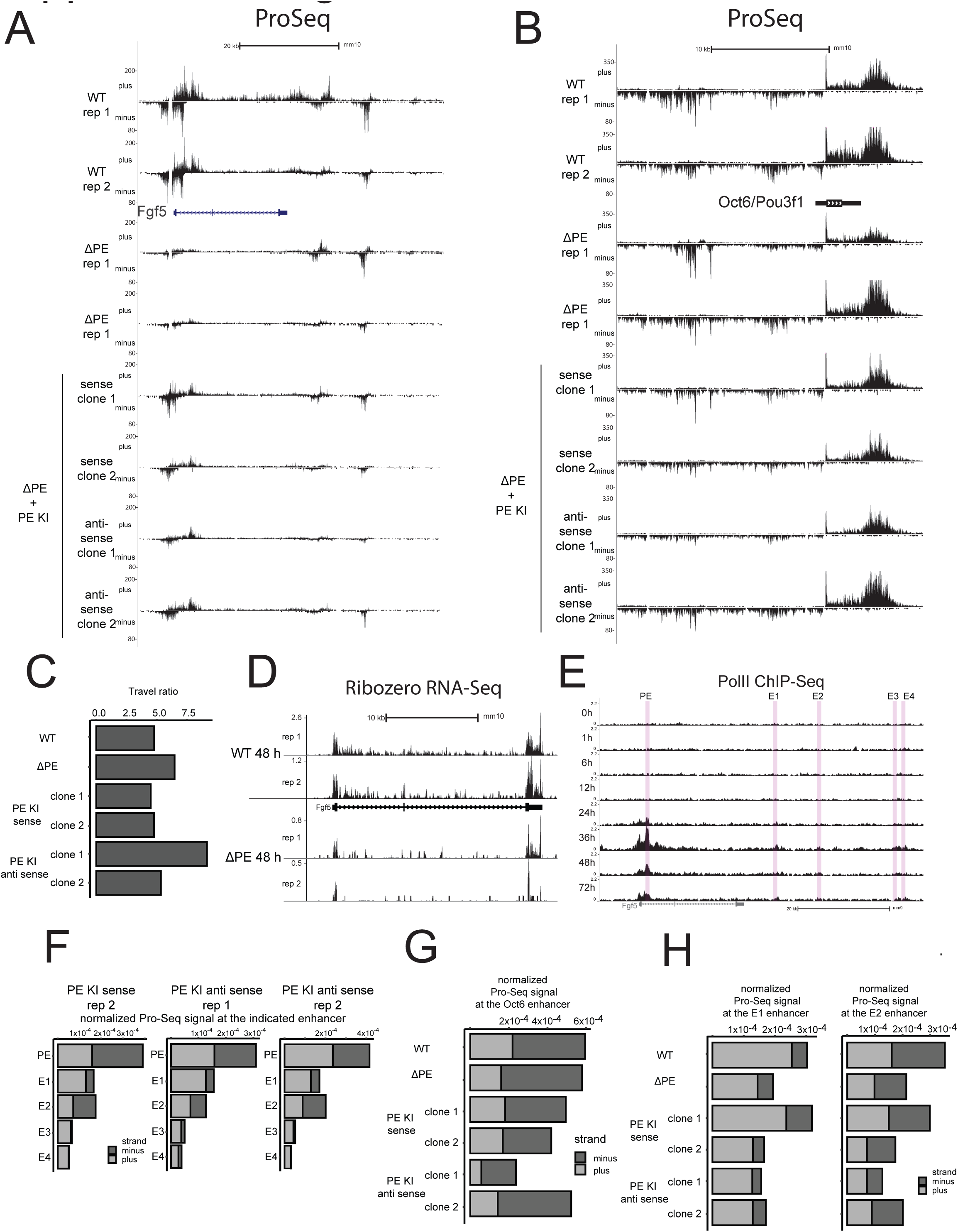
High levels of Pol II accumulate at the PE enhancer. (A) Spike-In-normalized strand-specific PRO-Seq signal at the *Fgf5* locus for WT and ΔPE cells (2 biological replicates each) as well as for all four PE KI clones after 40 h of ESC to EpiLC differentiation with fixed scale bar. (B) Spike-In-normalized strand-specific PRO-Seq signal at the *Oct6*/*Pou3f1* locus for WT and ΔPE cells (2 biological replicates each) as well as for all four PE KI clones after 40 h of ESC to EpiLC differentiation with fixed scale bar. (C) Travel ratio (PRO-Seq reads mapping on the plus strand between start of exon two and end of exon three divided by reads on the plus strand within a 350 bp window focused on the TSS) at the *Fgf5* gene for WT, ΔPE and PE KI cells after 40 h of ESC to EpiLC differentiation. (D) RiboZero RNA-Seq signal normalized for sequencing depth at the *Fgf5* locus in WT and ΔPE cells (2 biological replicates each) after 48 h of ESC to EpiLC differentiation with adjusted scale bar. (E) Pol II ChIP-Seq signal from Yang *et al*., 2019฀at the *Fgf5* locus along an ESC to EpiLC differentiation time course with fixed scale bar. Enhancers are highlighted in red. (F) Quantification of Spike-In-normalized PRO-Seq signal at the *Fgf5* enhancers on plus and minus strand in PE KI cells after 40 h of ESC to EpiLC differentiation. (G) Quantification of Spike-In-normalized PRO-Seq signal at the *Oct6* enhancer on plus and minus strand in WT, ΔPE and PE KI cells after 40 h of ESC to EpiLC differentiation. (H) Quantification of Spike-In-normalized PRO-Seq signal at the E1 and E2 *Fgf5* enhancers on plus and minus strand in WT, ΔPE and PE KI cells after 40 h of ESC to EpiLC differentiation.

